# Microrisk Lab: an online freeware for predictive microbiology

**DOI:** 10.1101/2020.07.23.218909

**Authors:** Yangtai Liu, Xiang Wang, Baolin Liu, Qingli Dong

## Abstract

Microrisk Lab was designed as an interactive modeling freeware to realize parameter estimation and model simulation in predictive microbiology. This tool was developed based on the R programming language and ‘Shinyapps.io’ server, and designed as a fully responsive interface to the internet-connected devices. A total of 36 peer-reviewed models were integrated for parameter estimation (including primary models of bacterial growth/ inactivation under static and non-isothermal conditions, secondary models of specific growth rate, and competition models of two-flora growth) and model simulation (including integrated models of deterministic or stochastic bacterial growth/ inactivation under static and non-isothermal conditions) in Microrisk Lab. Each modeling section was designed to provide numerical and graphical results with comprehensive statistical indicators depending on the appropriate dataset and/ or parameter setting. In this research, six case studies were reproduced in Microrisk Lab and compared in parallel to DMFit, GInaFiT, IPMP 2013/ GraphPad Prism, Bioinactivation FE, and @Risk, respectively. The estimated and simulated results demonstrated that the performance of Microrisk Lab was statistically equivalent to that of other existing modeling system in most cases. Microrisk Lab allowed for uniform user experience to implement microbial predictive modeling by its friendly interfaces, high-integration, and interconnectivity. It might become a useful tool for the microbial parameter determination and behavior simulation. Non-commercial users could freely access this application at https://microrisklab.shinyapps.io/english/.

## 1. Introduction

Foodborne pathogens have caused widespread food safety issues and potential severe risks nowadays (WHO, 2015). It is critical to understand and control the behavior (growth, survival or inactivation) or contaminated level of the focused microorganisms under different environmental conditions to ensure that foods are safe for consumption (Geeraerd, Valdramidis, & Van Impe, 2005; Augustin, 2011; González et al., 2018). For this reason, predictive microbiology has been developed as an efficient solution to estimate the bacterial concentration level in the perspective of mathematical modeling (Ross & McMeekin, 1994; Peleg & Corradini, 2011; Baranyi & Buss da Silva, 2017).

Microbiological predictive models are ordinarily classified as the primary model, secondary model, and tertiary model (Whiting & Buchanan, 1993). The primary model represents the relation between microbial concentrations and time under a specific condition by introducing the kinetic parameters, such as lag time, maximum specific growth/ inactivation rate, and decimal reduction time. While the secondary model describes the influence of environmental conditions on the kinetic parameters, such as growth and inactivation rates. The tertiary model refers to the computer program that integrates validated pertinent information to characterize the situation or explain the trend of the microbial contamination level under a specific condition (Whiting & Buchanan, 1993). Commonly, regression (or fitting) should be firstly applied to obtain the kinetic parameter and the effect of environmental conditions in accordance with the experimental observation (e.g. maximum population density, growth boundaries, and decimal reduction time). After identifying and validating the characteristic of the target microorganism(s), microbial behaviors (e.g. growth,inactivation, and survival) can be simulated under different conditions.

For realizing the parameter estimation, mathematical computing environments, such as R (www.R-project.org), MATLAB (The MathWorks, Inc., USA), and Python (www.python.org), are widely used in predictive microbiology. For example, ‘*nlsMicrobio*’ (Baty & Delignette-Muller, 2015) and ‘*Bioinactivation*’ (Garre, Fernández, Lindqvist, & Egea, 2017) are two packages dedicated to obtaining the microbial kinetic parameters in the R environment. However, the requirement of specific coding skills may increase the learning burden during the modeling process. Thus, many useful interactive modeling systems were developed in the last decades (Huang, 2014/2017b; Tenenhaus-Aziza & Ellouze, 2015; Dolan, Habtegebriel, Valdramidis & Mishra, 2015; Koutsoumanis, Lianou, & Gougouli, 2016). Among the developed freeware, IPMP 2013/ Global Fit (Huang, 2014/2017b), desktop DMFit, GInaFiT (Geeraerd, Valdramidis, & Van Impe, 2005/2006) and PMM-Lab (Plaza-Rodriguez et al., 2015) provided numerical and graphical interfaces for users to obtain different microbial model parameters without coding. These tools required to be installed and run under the desktop system of Windows or Mac OS. The online free tools, namely, the online DMFit of ComBase (www.combase.cc) and Bioinactivation FE (Garre et al, 2018) could be easily accessed via different internet-connected devices, which provided the ability of cross-platform to users.

On the other hand, some modeling systems put more emphasis on simulating or predicting the bacterial concentration level under different environmental conditions, which have some reference significance to microbial risk assessment and management. As the well-known free tools, Pathogen Modeling Program (USDA, 2016), and ComBase Predictor supported by their extensive microorganism-food database has been applied to predict the microbial behavior in culture medium or different food matrices. The applicability of a tertiary model is very dependent on the quantity and quality of the available knowledge integrated into the modeling system, such as experimental challenge test data, model types and associated model parameters. Recently, an updated application MicroHibro (González et al., 2018) allowed users to freely defined the model type and relevant parameter. This functionality may practically help users update the knowledge for the simulation when new evidence is observed. Meanwhile, it is also critical to take account of the uncertainty and variability of model parameters, especially in the application of the individual cell behavior modeling and risk assessment (Natau, 2001; Busschaert, Geeraerd, Uyttendaele, & Van Impe, 2011; Cornu et al., 2011; Koutsoumanis & Lianou, 2013; Alonso, Molina, & Theodoropoulos, 2014; Augustin et al., 2014). Thus, it is essential to introduce the stochastic approach in the prediction and simulation study.

Besides, much more complex situations should be considered to describe the microbial behavior in the real food chain, namely, the coexistence of multi-microorganisms, and the concentration change under dynamic conditions (Iannetti et al., 2017; Li, Huang, & Yuan, 2017, Göransson, Nilsson, & Jevinger, 2018; Ndraha et al, 2018; Hwang & Huang, 2018). In non-isothermal modeling, free tools of ComBase Predictor, IPMP Dynamic Prediction (USDA, 2017), GroPIN (https://www.aua.gr/psomas/gropin/), FSSP (http://fssp.food.dtu.dk), and UGPM (Psomas, Nychas, Haroutounian, & Skandamis, 2011) were designed for microbial simulation. The web-based tool, Bioinactivion FE, was recently developed for fitting and simulating microbial inactivation under isothermal or non-isothermal conditions (Garre et al, 2018). This tool exactly facilitated scientists handle different inactivation analyses without the need to code the mathematical models in a programming environment. However, there was still a lack of tools for kinetic analysis on the microbial dynamic growth (Tenenhaus-Aziza & Ellouze, 2015; Koutsoumanis, Lianou, & Gougouli, 2016). Hence, it may be helpful to design an integrated system containing the functionality for parameter estimation and model simulation under non-isothermal conditions.

This research introduced the main features of Microrisk Lab, an online modeling system integrating comprehensive microbial predictive models. Six case studies were implemented to describe a part of functionality and performance of this new application for parameter estimation and model simulation in predictive microbiology. The first version of Microrisk Lab was deployed on the ‘Shinyapps.io’ server, and available at https://microrisklab.shinyapps.io/english/ (in English) and https://microrisklab.shinyapps.io/chinese/ (in Chinese).

## 2. Materials and methods

### 2.1. Design logic and programming basics of Microrisk Lab

Microrisk Lab was designed as a R-based web application with a user-friendly interface for performing parameter estimation or model simulation studies in predictive microbiology. The coding language R, an open-source mathematical environment, could run on a wide variety of computer systems, including Windows, UNIX, and Mac OS. Several basic R packages, such as ‘*ggplot2’* (Wickham et al., 2019), ‘*mc2d’* (Pouillot & Delignette-Muller, 2010), and ‘*Metrics’* (Hamner, B., Frasco, M., & LeDell, E., 2018), were referenced in this tool for mathematical and statistical analysis (see supplementary data). Meanwhile, the platform of ‘*Shiny*’ (http://shiny.rstudio.com/), *shinydashboard’* (Chang & Borges Ribeiro, 2019), and ‘*plotly*’ (https://plot.ly) were introduced to improve the operability and practicability of Microrisk Lab. The simple graphical user interface (GUI) and interactive output can automatically adapt to different screen sizes (Fig.1). Each section has a uniform interactive logic from left to right (horizontal view) or up to down (vertical view) corresponding to problem selection, condition setting, and result analysis. The observed measurement for parameter estimation or model simulation can be directly typed in the data dialog or pasted from other table files. After submitting all condition settings, users are free to make a real-time control on the interactive plot for better visualization then save as the local image file (Portable Network Frame file).

**Fig. 1.**
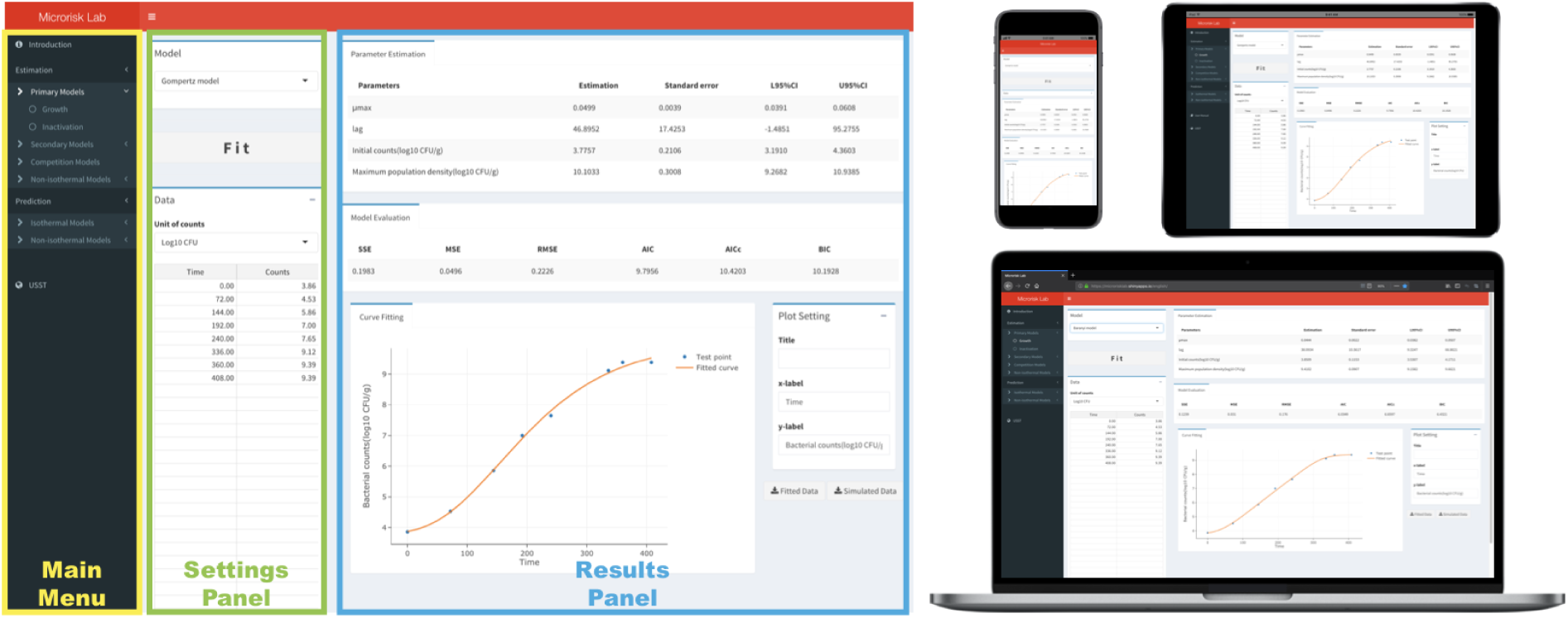
Overview of the layout of Microrisk Lab and its visual interface on different internet-connected devices.

The structural framework of Microrisk lab is shown in Fig.2, which is basically divided into the ‘Estimation’ and ‘Simulation’ module. The ‘Estimation’ module was focused on determining the microbial parameters by the experimental observations under different conditions. The ‘Simulation’ module aimed to simulate the bacterial concentration changes under different temperatures by using different built-in predictive models.

**Fig. 2.**
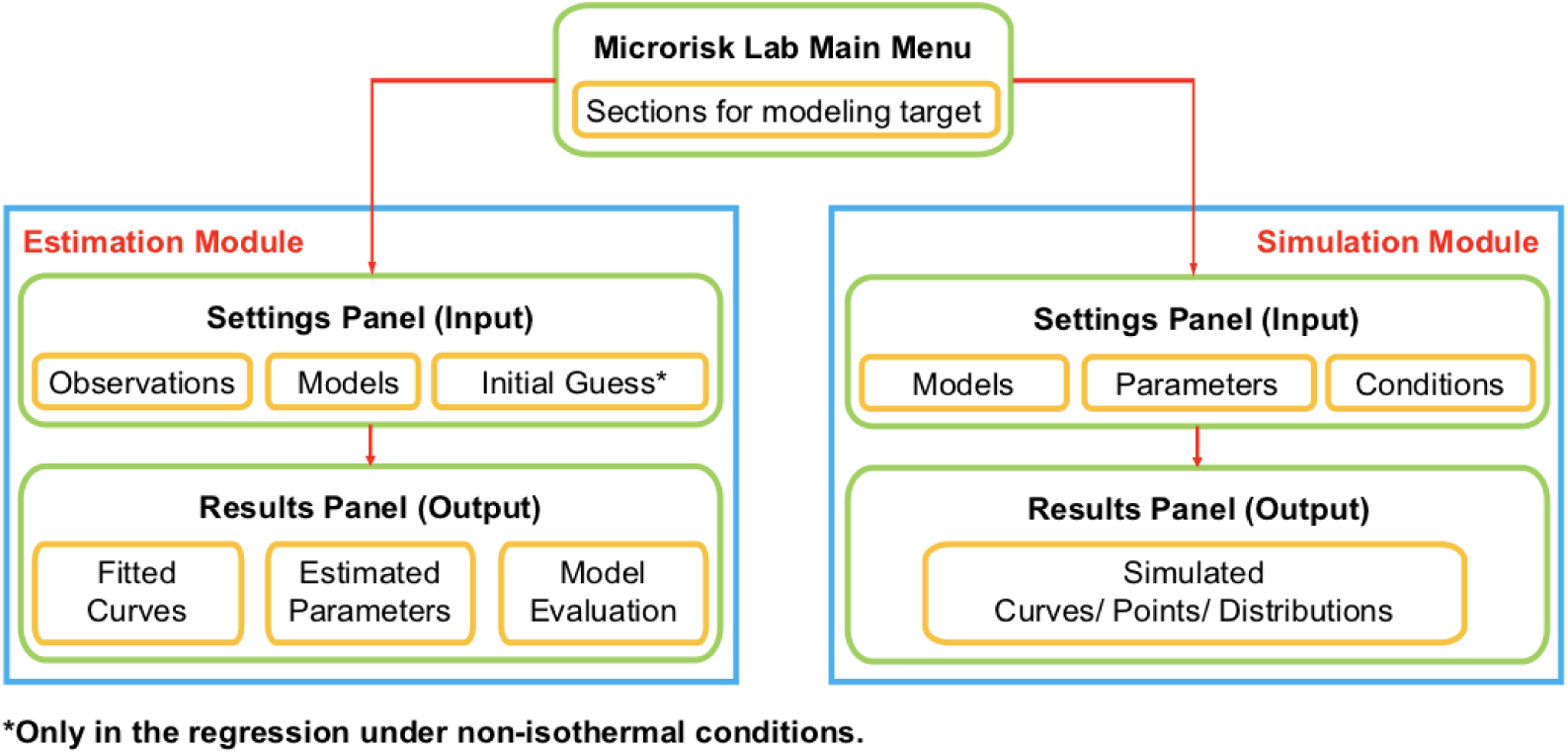
The structural framework and workflow of Microrisk Lab.

In the ‘Estimation’ module, the least-squares method was implemented to search the optimized model parameter, which was conducted by the ***nls*** function in the ‘*stats*’ package. Both ‘NL2SOL’ algorithm (for the dynamic regression) and Gauss-Newton algorithm (for other regressions) were used in Microrisk Lab. If the fitting is successful, results of the fitted curve, parameter estimation, and model evaluation should be reported in the ‘Results Panel’. Meanwhile, the raw and generated datasets (observed, fitted, and simulated data) are downloadable as ‘csv.’ files. Otherwise, a pop-up window would remind the user that regression is failure.

The ‘Simulation’ module in Microrisk Lab does not restrict the type of microorganisms or food. The microbial growth and inactivation should be simulated by defining the model type, microbial kinetic parameter, and temperature condition (or time-temperature profile). Besides, the stochastic simulation can be performed at static conditions. In this case, probability distribution of the parameter and condition are defined according to the mean value and standard deviation. Here, the duration of growth or inactivation steps is assumed as a Uniform distribution, and other default parameter settings are assumed as the Normal distribution. According to former researches (Baranyi, George, & Kutalik, 2009; Koutsoumanis & Lianou, 2013; Huang 2016), the LogNormal/ Gamma distribution and LogNormal/ Logistic distribution were additionally considered in the parameter setting of lag time (shoulder) and specific growth rate, respectively. Then the stochastic model can be conducted by using the simple sampling method with optional 100/1,000/10,000 iteration times for Monte-Carlo simulation.

### 2.2. Mathematical models and statistical indicators in Microrisk Lab

In version 1.0, Microrisk Lab contained 36 peer-reviewed models to implement parameter estimation or model simulation in predictive microbiology. Specifically, 20 explicit equations were chosen by considering different shapes of the growth/ inactivation curve for microbial dynamics under static conditions (Tab.1); 10 secondary models were selected in view of the impact of temperature/ pH/ water activity on the specific growth rate (Tab.2); 2 piecewise functions were applied to describe two flora competition growth (Tab.3); and 4 groups of ordinary differential equations were presented by combining the primary model and secondary model for microbial growth/ inactivation under non-isothermal conditions (Tab.3). The definition of each parameter was illustrated in the list of symbols.

**Table 1.**
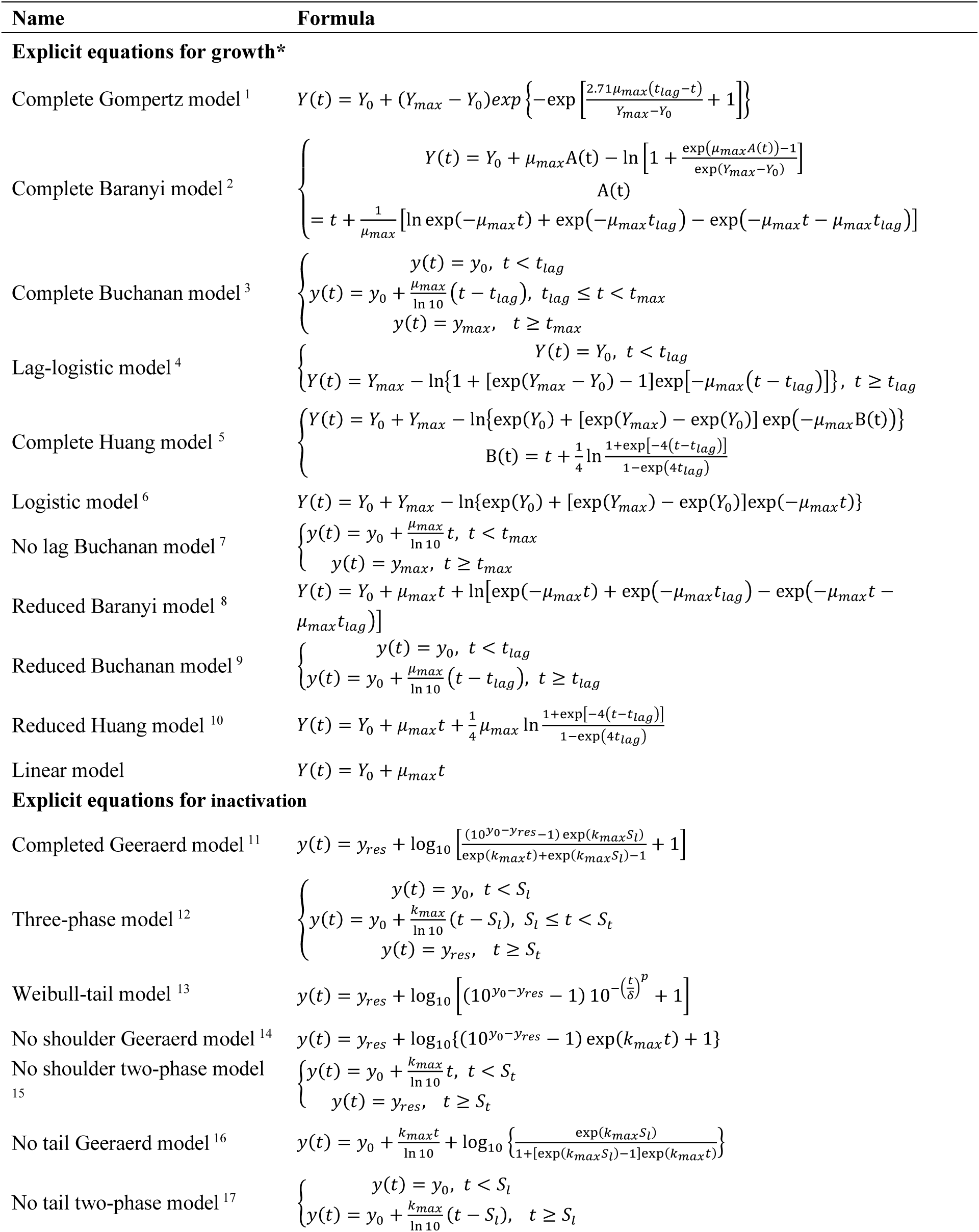
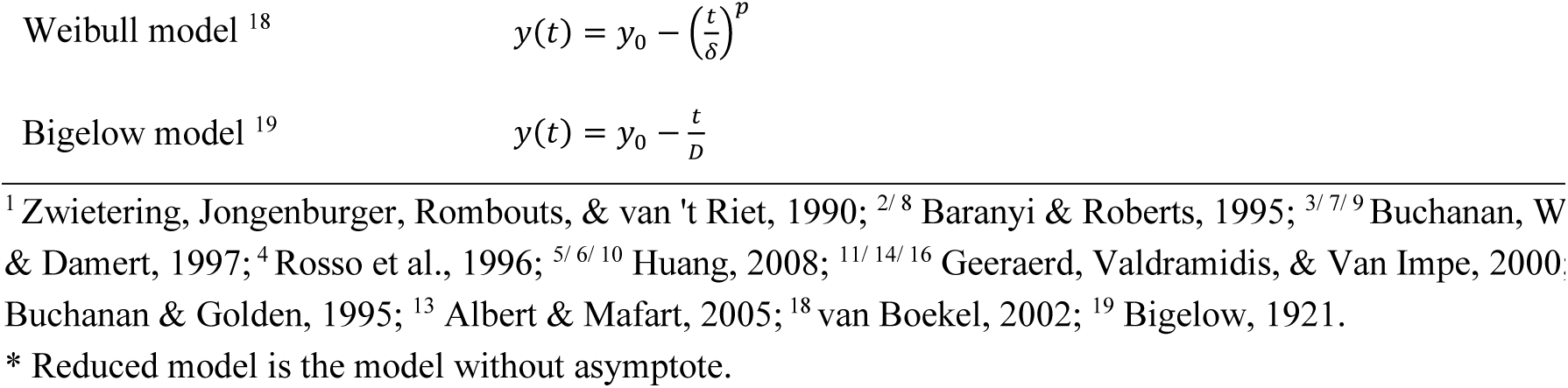
Primary models included in microrisk Lab

**Table 2.**
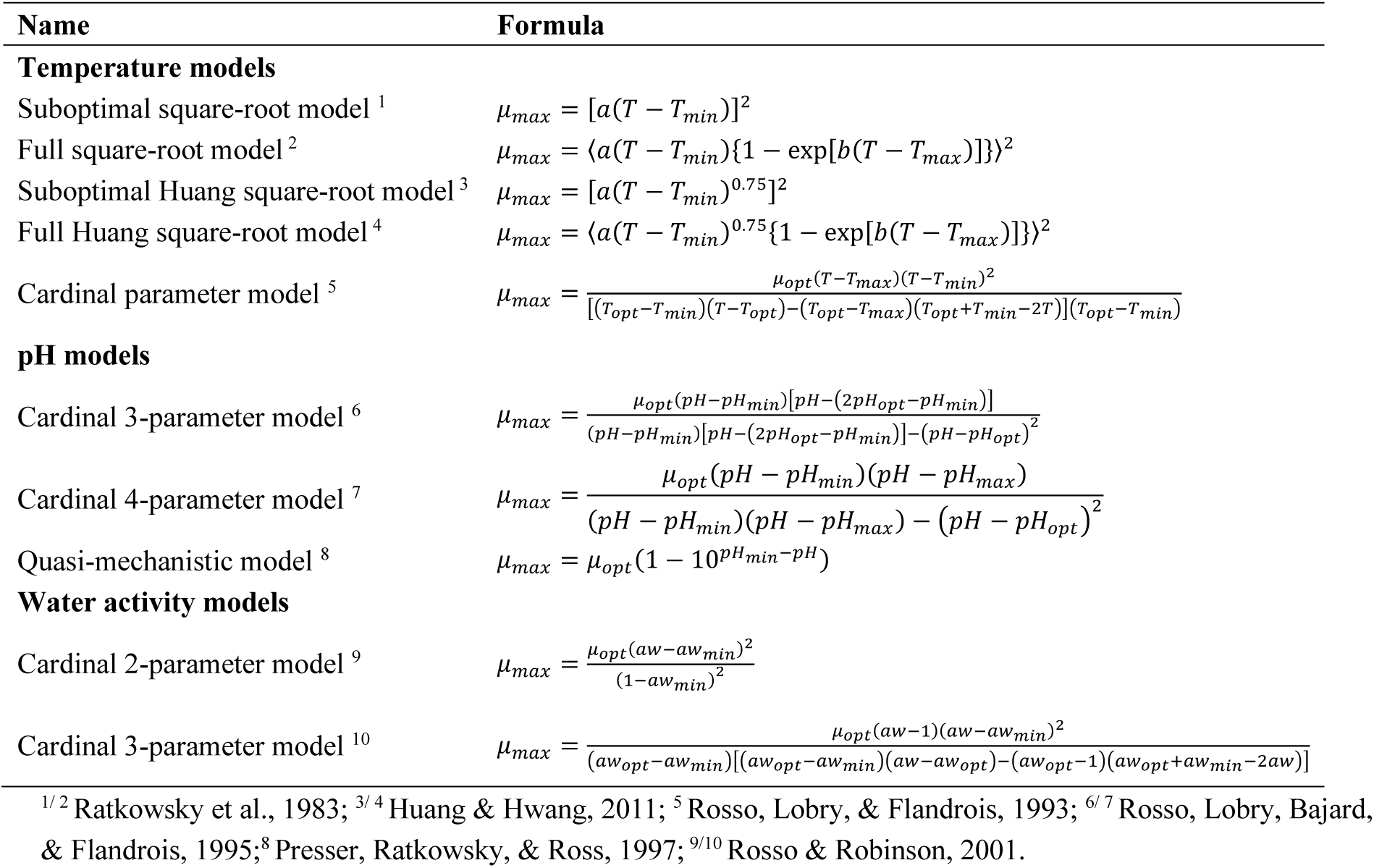
Secondary models for *μ*_*max*_ included in Microrisk Lab

**Table 3.**
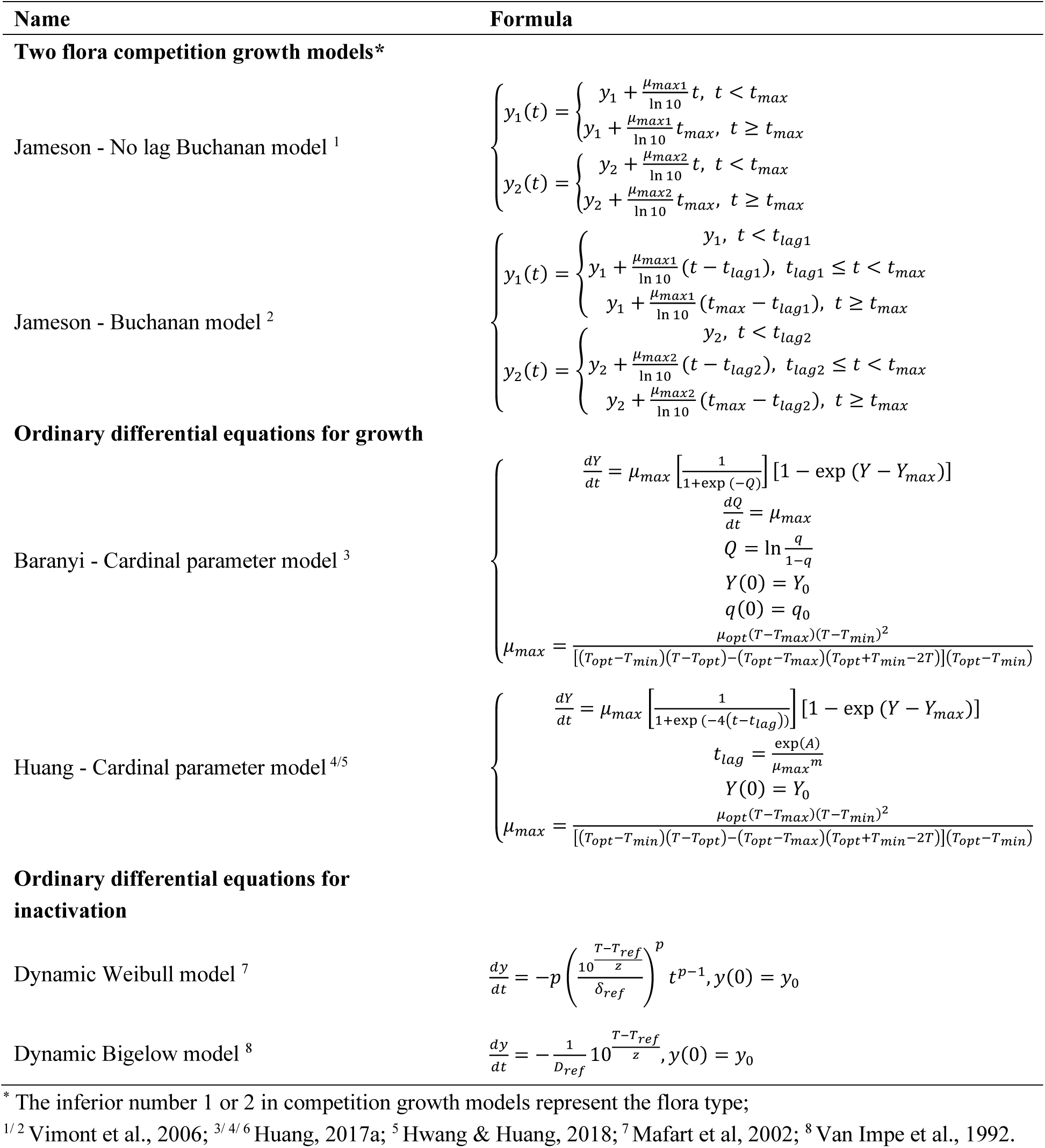
Complex models included in Microrisk Lab

| Name | Formula |
| --- | --- |
| **Two flora competition growth models\*** |  |
| Jameson - No lag Buchanan model <sup>1</sup> | $\begin{cases} y_1(t) = \begin{cases} y_1 + \frac{\mu_{max1}}{\ln 10} t, & t < t_{max} \\ y_1 + \frac{\mu_{max1}}{\ln 10} t_{max}, & t \geq t_{max} \end{cases} \\ y_2(t) = \begin{cases} y_2 + \frac{\mu_{max2}}{\ln 10} t, & t < t_{max} \\ y_2 + \frac{\mu_{max2}}{\ln 10} t_{max}, & t \geq t_{max} \end{cases} \end{cases}$ |
| Jameson - Buchanan model <sup>2</sup> | $\begin{cases} y_1(t) = \begin{cases} y_1, & t < t_{lag1} \\ y_1 + \frac{\mu_{max1}}{\ln 10} (t - t_{lag1}), & t_{lag1} \leq t < t_{max} \\ y_1 + \frac{\mu_{max1}}{\ln 10} (t_{max} - t_{lag1}), & t \geq t_{max} \end{cases} \\ y_2(t) = \begin{cases} y_2, & t < t_{lag2} \\ y_2 + \frac{\mu_{max2}}{\ln 10} (t - t_{lag2}), & t_{lag2} \leq t < t_{max} \\ y_2 + \frac{\mu_{max2}}{\ln 10} (t_{max} - t_{lag2}), & t \geq t_{max} \end{cases} \end{cases}$ |
| **Ordinary differential equations for growth** |  |
| Baranyi - Cardinal parameter model <sup>3</sup> | $\begin{cases} \frac{dY}{dt} = \mu_{max} \left[ \frac{1}{1 + \exp(-Q)} \right] [1 - \exp(Y - Y_{max})] \\ \frac{dQ}{dt} = \mu_{max} \\ Q = \ln \frac{q}{1-q} \\ Y(0) = Y_0 \\ q(0) = q_0 \\ \mu_{max} = \frac{\mu_{opt}(T - T_{max})(T - T_{min})^2}{[(T_{opt} - T_{min})(T - T_{opt}) - (T_{opt} - T_{max})(T_{opt} + T_{min} - 2T)](T_{opt} - T_{min})} \end{cases}$ |
| Huang - Cardinal parameter model <sup>4/5</sup> | $\begin{cases} \frac{dY}{dt} = \mu_{max} \left[ \frac{1}{1 + \exp(-4(t - t_{lag}))} \right] [1 - \exp(Y - Y_{max})] \\ t_{lag} = \frac{\exp(A)}{\mu_{max}^m} \\ Y(0) = Y_0 \\ \mu_{max} = \frac{\mu_{opt}(T - T_{max})(T - T_{min})^2}{[(T_{opt} - T_{min})(T - T_{opt}) - (T_{opt} - T_{max})(T_{opt} + T_{min} - 2T)](T_{opt} - T_{min})} \end{cases}$ |
| **Ordinary differential equations for inactivation** |  |
| Dynamic Weibull model <sup>7</sup> | $\frac{dy}{dt} = -p \left( \frac{10^{\frac{T - T_{ref}}{z}}}{\delta_{ref}} \right)^p t^{p-1}, y(0) = y_0$ |
| Dynamic Bigelow model <sup>8</sup> | $\frac{dy}{dt} = -\frac{1}{D_{ref}} 10^{\frac{T - T_{ref}}{z}}, y(0) = y_0$ |

\* The inferior number 1 or 2 in competition growth models represent the flora type;
<sup>1/2</sup> Vimont et al., 2006; <sup>3/4/6</sup> Huang, 2017a; <sup>5</sup> Hwang & Huang, 2018; <sup>7</sup> Mafart et al, 2002; <sup>8</sup> Van Impe et al., 1992.

Note that the 2^nd^ order Runge-Kutta method or Heun’s method (Eq.1, Press, Teukolsky, Vetterling, & Flannery, 2007) was applied as the rapid numerical method to solve the ordinary differential equations in the dynamic kinetic analysis. During the computational procedures, the non-isothermal growth/ inactivation was firstly solved by the 2^nd^ order Runge-Kutta method to calculate the predicted value, corresponding to each of the sampling time for bacterial counting. Then, the predicted values were applied to match the observed values by a nonlinear least-squares function to determine the optimized parameter estimation. Similar algorithm of the 4^th^ order Runge-Kutta method was also realized by R and other programming languages in previous studies (Press, Teukolsky, Vetterling, & Flannery, 2007; Cattani et al., 2016; Li et al., 2017; Huang, 2017a; Hwang & Huang, 2018). The time step (0.1, 0.01, or 0.001) could be selected by the user in the regression of non-isothermal growth and inactivation.

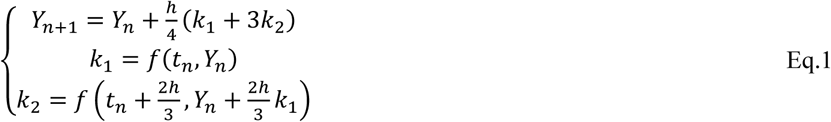

In the module of parameter estimation, a recognition algorithm (if/ else statement) was preset to transfer the input (counting) data into the appropriate unite before fitting to a specific model, which allowed users to freely choose the preferred input unit of the counting data (”Log10 CFU/g or CFU/ml”, “Ln CFU/g or CFU/ml”, or “CFU/g or CFU/ml”) in Microrisk Lab. Meanwhile, results of the model parameter, the estimated value, standard error, and lower and upper 95% confidence intervals (Eq.2), were provided by the R package of *“stats”* and *“nlstool”*. After obtaining the estimated and evaluated values, users could select the decimal digits (0, 1, 2, 3, or 4) of the generated results, which should be determined according to the unit precision of the parameter.

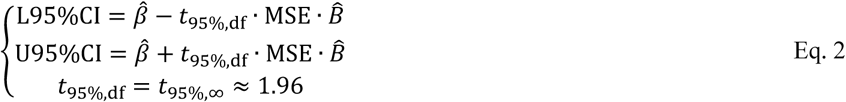

where 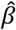 is the estimated parameter; MSE is the mean sum of square error; 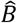 is the inverse of the matrix of second derivatives of the log-likelihood function as a function of *β* evaluated at the parameter estimates 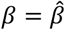; df is degrees of freedom, which is assumed infinite; *t*_95%,df_ is the value from the t distribution for 95% confidence for the specified number of df.

Furthermore, several statistical indicators were reported to evaluate and compare the goodness-of-fit between observed and predicted values, such as the residual sum of squares (RSS, Eq.3, Draper & Smith, 1998), mean sum of squared error (MSE, Eq.4, Geeraerd et al., 2005), root mean sum of squared error (RMSE, Eq.5, Ratkowsky, 2003), regular Akaike information criterion (AIC, Eq.6, Akaike, 1974), corrected AIC (AICc, Eq.7, Burnham & Anderson, 2003) and Bayesian information criterions (BIC, Eq.8, Schwarz, 1978). As pointed out by Ratkowsky (2003), the coefficient of determination (R^2^, Eq.9, Rawlings, Pantula, & Dickey, 2001) and the adjusted coefficient of determination (Adjusted R^2^, Eq.10, Rawlings, Pantula, & Dickey, 2001) might be inappropriate to evaluate the non-linear models. Thus, Microrisk Lab provided these two indicators only for linear models.

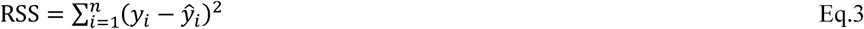

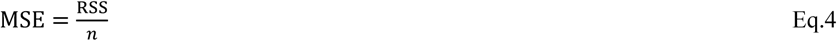

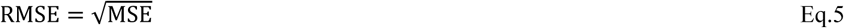

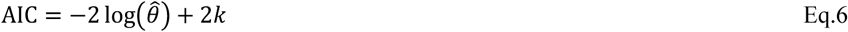

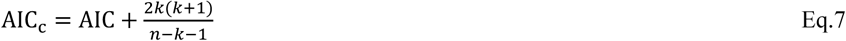

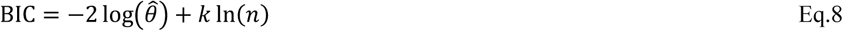

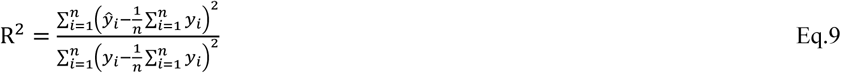

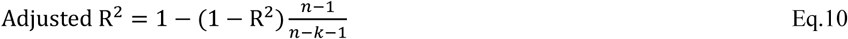

where *y*_*i*_ is the i th value of the observation; 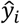 is the i th value of the prediction; *k* is the number of parameters; and *n* is the number of sample data; 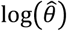 is the numerical value of the log-likelihood for the fitted model (the probability of the data given a model in the model), which is donated by the **logLik()** function built in the R package *‘stats’*.

Besides, for stochastic simulation, the Pearson correlation coefficient (Eq.11) is also calculated to measure the linear correlation between different model variables (***P***) and the final bacterial concentration (*y*_*final*_). The dependence or association relationship can be measured by the generated tornado plot.

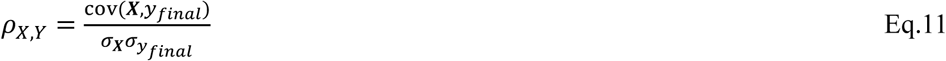

where cov(***X***, *y*_*final*_) is the covariance of the final bacterial concentration and different model variables; *σ*_***X***_ is the standard deviation of different model variables; 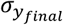 is the standard deviation of the final bacterial concentration.

### 2.3. Practical examples for Microrisk Lab

To illustrate the performance of Microrisk Lab, we collected 6 datasets from the peer-reviewed papers and lab observation for parameter estimation and simulation. Specifically, the study on the static/ non-isothermal growth regression, static/ non-isothermal inactivation regression, secondary model regression, and static stochastic growth simulation. The datasets for the kinetic analyses (Case I – V) were attached in the supplementary data. It should be noted that only a part of models was compared with the relevant modeling system in this study. More results on the comparison between built-in models were provided in the user manual (see supplementary data).

#### 2.3.1. Case I – Kinetic analysis of *Listeria monocytogenes/ Listeria innocua* growth under a static condition

A growth measurement of *L. monocytogenes*/ *L. innocua* in tryptose phosphate broth (TPB) was obtained from the ComBase browser (ComBase ID: LM127_11) according to the research of Buchanan & Phillips (1990). In order to compare with the online DMFit and Excel DMFit, the ‘Complete Baranyi model’ in Microrisk Lab was chosen to determine the kinetic parameter of *L. monocytogenes*.

#### 2.3.2. Case II – Kinetic analysis of *Salmonella enterica* inactivation under a static condition

A thermal inactivation curve of *S. enterica* in Brain Heart Infusion (BHI) under 60°C reported by Wang, Devlieghere, Geeraerd, & Uyttendaele (2017) was used to evaluate the inactivation model in Microrisk Lab. According to the suggestion by the author, ‘Log-linear + Shoulder’ model in GInaFiT (version 1.7) was selected for fitting. Therefore, performance of ‘No tail Geeraerd model’ in Microrisk Lab was compared in parallel with GInaFiT as well.

#### 2.3.3. Case III– Effect of temperature on the specific growth rate of *Salmonella* Typhimurium

We cited a study on the maximum specific growth rate of *S.* Typhimurium (ATCC 14028) in chicken breast (Oscar, 2002) to estimate the growth boundary and optimal parameter by fitting the cardinal parameters model. The value of the specific growth rate under different static temperature conditions was converted to the same units (natural logarithm) in Microrisk Lab before regression. Both IPMP 2013 and Prism (version 7.0, GraphPad Software, USA) were applied for comparison.

#### 2.3.4. Case IV – Kinetic analysis of *L. monocytogenes* growth under non-isothermal conditions

For growth modeling under non-isothermal conditions, the observed concentration and time-temperature profile were introduced from a study on *L. monocytogenes* growth in cooked beef samples under non-isothermal conditions. During the experiments, four *L. monocytogenes* strains (serotype 1/2a, 1/2b, 1/2c and 4b, meat isolated) were inoculated in a heat-treated ready-to-eat braised beef product (ca. 1% NaCl, pH=6.2, aw=0.983) and incubated in an air-packaged sterile stomacher bag under the fluctuating temperature ranging from 5 to 40°C. To date, there were no other integrated systems specialized for non-isothermal growth regression analysis. Thus, the measurements would be fitted by the ‘Baranyi-Cardinal parameter model’ and ‘Huang-Cardinal parameter model’ in Microrisk Lab.

#### 2.3.5. Case V – Kinetic analysis of *Bacillus sporothermodurans* IC4 spores inactivation under non-isothermal conditions

In this case, a dataset was adopted from the supplementary data of the verification research on the non-isothermal inactivation modeling by Bioinactivation core (Garre, Fernández, Lindqvist, & Egea, 2017). This example data described the inactivation of *B. sporothermodurans* IC4 spores under non-isothermal heating conditions. Bioinactivation FE (Garre et al, 2018), a web tool based on Bioinactivation core, was introduced to compare for the estimated results with Microrisk Lab. The dynamic Bigelow model was selected with the non-linear regression algorithm for inactivation fitting under non-isothermal conditions.

#### 2.3.6. Case VI – Simulation of *S.* Typhimurium stochastic growth under a static condition

The stochastic simulation was based on the study of Koutsoumanis & Lianou (2013) which obtained the growth parameters of *S.* Typhimurium individual cells with an automated time-lapse microscopy method. A 10,000 times Monte-Carlo simulation was realized in commercial software, @Risk for Excel (version 6.0, Palisade Corporation, USA), to describe the stochastic growth of *S.* Typhimurium individual cells. According to the distribution of the conditions and parameters, the stochastic growth of a single cell with the Buchanan model was reproduced in Microrisk Lab for comparison.

## 3. Results and discussion

### 3.1. Comparison of the primary and secondary modeling

Case studies of the growth/ static inactivation under static conditions and the effect of temperature on the specific growth rate were evaluated in Microrisk Lab and compared with other integrated modeling systems. The fitted curves of Case I, Case II, and Case III downloaded from Microrisk Lab are shown in Fig.3, which illustrates the consistency in the result rendering of different sections. Note that the interactivity of Microrisk Lab allows users to change the coordinate axis settings and the displayed results freely.

**Fig. 3.**
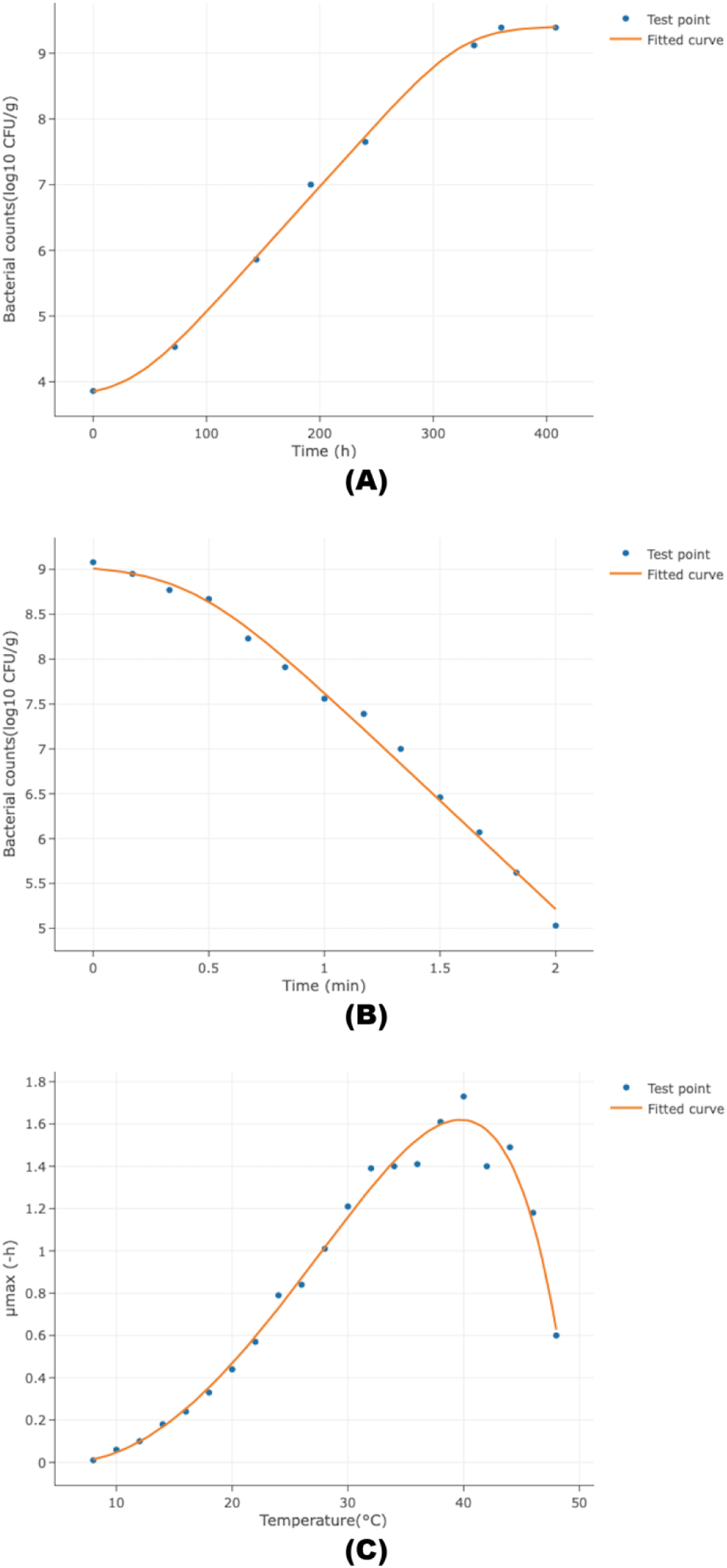
The fitted curve of (A) Case I with the ‘Complete Baranyi model’, (B) Case II with the ‘No tail Geeraerd model’, and (C) Case III with the ‘Cardinal parameter model’ downloaded from Microrisk Lab. (The blue dot represents the observed bacterial count, and the origin line represents the fitted curve.)

Tab.4 lists the results of the estimation and evaluation in Microrisk Lab and DMFit by fitting the complete Baranyi model for Case I. Although most of the estimated results were similar, there was around four-hour distinction between Online DMFit and Microrisk Lab/ Excel DMFit on the estimated lag time. It should be noted that, in the DMFit systems, two curvature parameters of model need to be determined and fixed before regression. According to the help documentation for Online DMFit (https://browser.combase.cc/DMFit_Help.aspx) and manual for Excel DMFit (version 3.5), the default values for two curvature parameters, nCurv and mCurv, were 1 and 10, respectively. In contrast, all estimable parameters were determined by globally searching for the optimized estimates in Microrisk Lab, which could also cause the discrepancy of results. The evaluation indicators and standard errors of parameters are getting close to that in Microrisk Lab when increasing the value of nCurv from default 1 to 2 in Excel DMFit. However, it is noticeable that the reason for differences of the estimated value between the online DMFit and Excel DMFit is inexplicable. Meanwhile, the model evaluation indicators were different in DMFit tools and Microrisk Lab, we further calculated adjusted R^2^ by Eq.8 according to the regression in Microrisk Lab for comparison (Tab.4). The results illustrate that the estimated adjusted R^2^ has no obvious differences between Microrisk Lab and DMFit tools with different curvature settings.

**Table 4.** Comparison on static growth fitting results of Microrisk Lab and DMFit (Complete Baranyi model)

|  | Microrisk Lab | Online DMFit | Excel DMFit |  |  |
| --- | --- | --- | --- | --- | --- |
| Curvature paramters | - | Default | nCurv=1 (Default) | nCurv=1.5 | nCurv=2 |
| **Parameter estimation** |  |  |  |  |  |
| Parameters | Est. (SE)*<br>(95% CI) ** | Est. (SE)* | Est. (SE)* | Est. (SE)* | Est. (SE)* |
| $y_0$ (log10 CFU/g) | 3.85 (0.12)<br>(3.53, 4.17) | 3.84 (0.12) | 3.82 (-) | 3.82 (-) | 3.86 (-) |
| $y_{max}$ (log10 CFU/g) | 9.41 (0.09)<br>(9.16, 9.66) | 9.44 (0.10) | 9.46 (0.12) | 9.46 (0.11) | 9.46 (0.10) |
| $t_{lag}$ (h) | 38.09 (10.36)<br>(9.32, 66.86) | 42.72 (11.35) | 38.89 (16.08) | 38.66 (11.69) | 38.01 (10.10) |
| $\mu_{max}$ (1/h) | 0.044 (0.002)<br>(0.038, 0.051) | 0.046 (0.003) | 0.045 (0.004) | 0.045 (0.004) | 0.044 (0.002) |
| **Model evaluation** |  |  |  |  |  |
| RSS | 0.1239 | - | - | - | - |
| MSE | 0.0310 | - | - | - | - |
| RMSE | 0.1760 | - | - | - | - |
| AIC | 4.0349 | - | - | - | - |
| AICc | 9.3683 | - | - | - | - |
| BIC | 4.3527 | - | - | - | - |
| Adjusted R<sup>2</sup> | 0.9976*** | 0.997 | 0.9970 | 0.9974 | 0.9975 |

\* Est.: Estimation; SE: Standard error.
\*\* 95%CI: lower and upper 95% confidence intervals.
\*\*\* Results no show in Microrisk Lab

As listed in Tab.5, for the static inactivation modeling, results of estimated parameters and evaluation indicators show no difference between Microrisk Lab and GInaFiT 1.7 when using the same model. Similarly, the effect of temperature on the *μ*_*max*_ of *S.* Typhimurium in chicken breast has been equivalently described in Microrisk Lab, IPMP 2013, and GraphPad Prism by the cardinal parameters model (Tab.6). Remember that the equation of AIC built-in IPMP 2013 was referred to the study by van Boekel, & Zwietering (2007), which was different from that of built-in Microrisk Lab. Above results indicated that Microrisk Lab could offer an equivalent accuracy to other integrated systems on primary and secondary modeling studies.

**Table 5.**
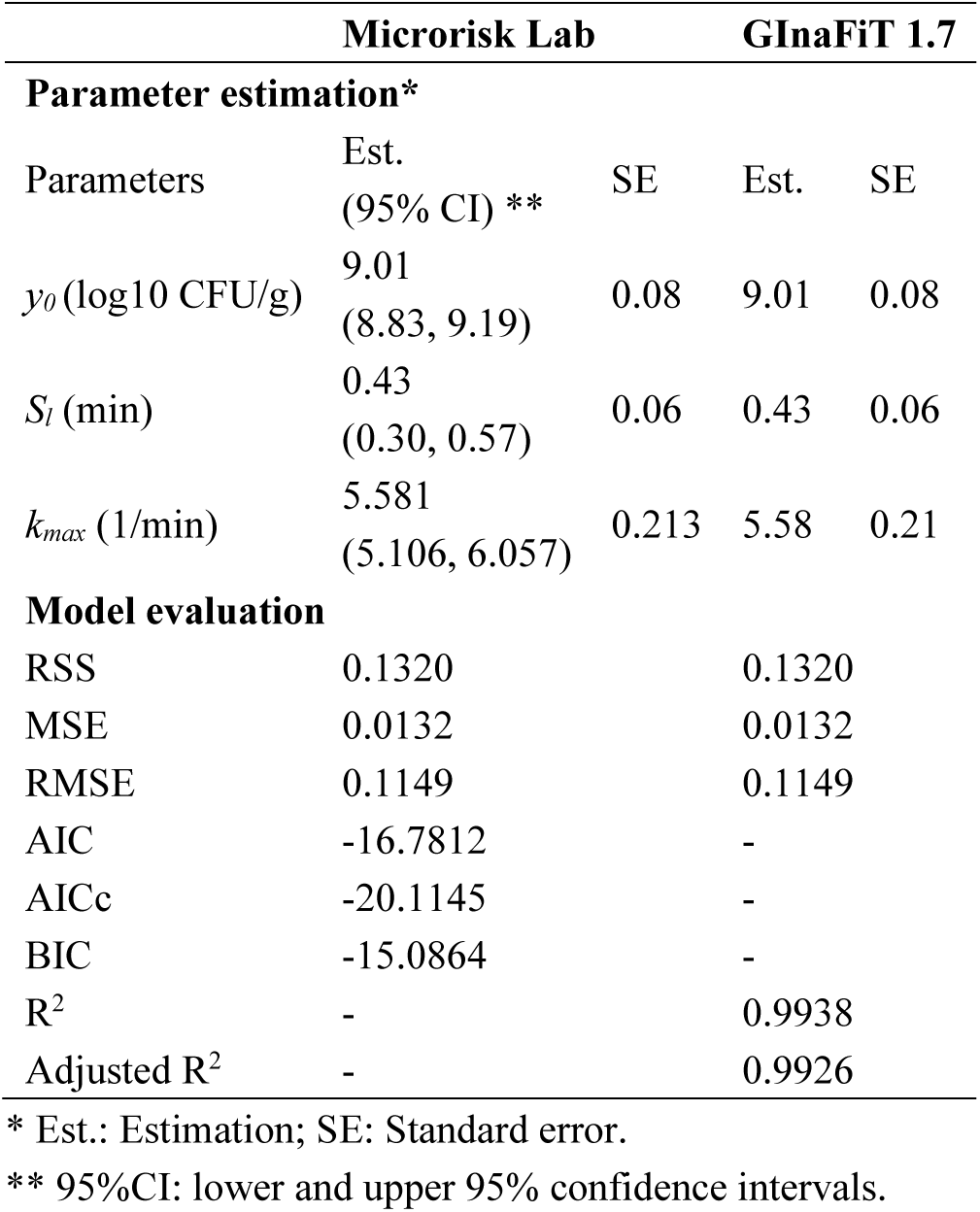
Comparison on inactivation fitting results of Microrisk Lab and GInaFiT (No tail Geeraerd model)

**Table 6.**
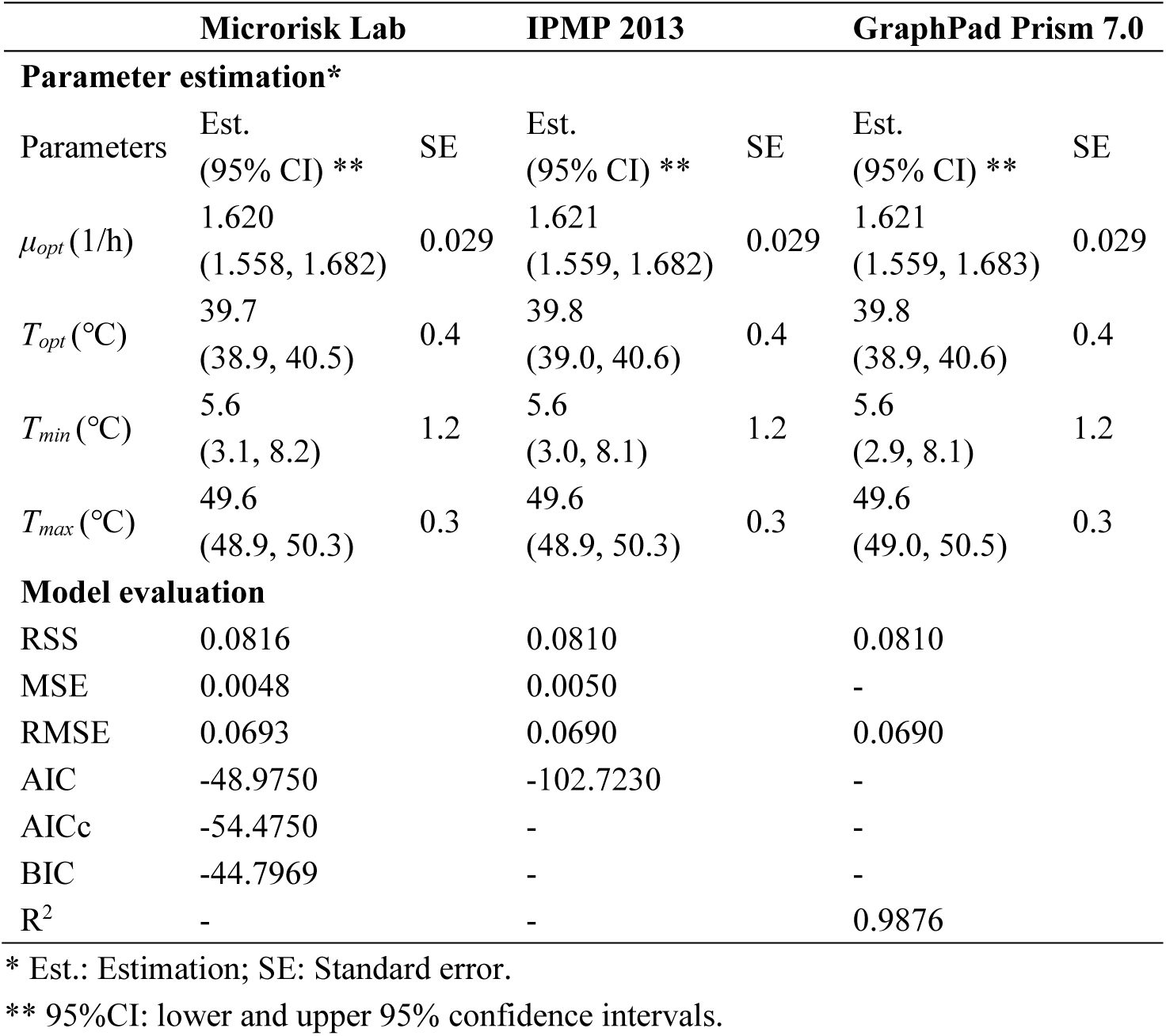
Comparison on secondary model fitting results of Microrisk Lab and IPMP 2013 (Cardinal parameter model)

### 3.2. Comparison of the dynamic modeling

In Case IV, both time-temperature profile and bacterial counting data were needed for the dynamic analysis. Initial guesses of the model parameter were required to assist in regression converge easily. According to former studies (ICMSF, 1996; Magalhães et al., 2014), *L. monocytogenes* probably has a growth temperature range from 0 to 45°C, the optimal specific growth rate is around 1ln CFU/h (or 1/h) under 37°C in meat products. Initial guesses (Default values) of *q*_0_, *A*, and *m* are preset as 1 in Microrisk Lab when there has no reliable basic knowledge on these parameters. With the above initial settings, both regressions could converge successfully. The fitted curve and the estimated result are exhibited in Fig.4 and Tab.7, respectively. The results illustrated that the microbial growth parameters could be obtained from Microrisk Lab with the measurements under non-isothermal conditions in one analysis. Meanwhile, the Baranyi - Cardinal parameter model and Huang - Cardinal parameter model could well describe the non-isothermal growth of *L. monocytogenes* in cooked beef.

**Table 7.** Non-isothermal growth model fitting results of Microrisk Lab

| Baranyi-Cardinal parameter model |  |  |  | Huang-Cardinal parameter model |  |  |  |
| --- | --- | --- | --- | --- | --- | --- | --- |
| Parameter estimation* |  |  |  |  |  |  |  |
| Parameters | Int. | Est.<br>(95% CI) ** | SE | Parameters | Int. | Est.<br>(95% CI) ** | SE |
| $y_0$ (log10 CFU/g) | - | 3.39<br>(3.36, 3.43) | 0.02 | $y_0$ (log10 CFU/g) | - | 3.45<br>(3.41, 3.50) | 0.02 |
| $y_{max}$ (log10 CFU/g) | - | 8.21<br>(8.18, 8.25) | 0.02 | $y_{max}$ (log10 CFU/g) | - | 8.21<br>(8.16, 8.27) | 0.03 |
| $\mu_{opt}$ (1/h) | 1.000 | 1.065<br>(0.854, 1.276) | 0.096 | $\mu_{opt}$ (-h) | 1.000 | 1.242<br>(0.825, 1.659) | 0.187 |
| $T_{opt}$ (°C) | 37.0 | 36.4<br>(35.4, 37.5) | 0.5 | $T_{opt}$ (°C) | 37.0 | 38.0<br>(33.5, 42.4) | 2.0 |
| $T_{min}$ (°C) | 0.0 | -1.1<br>(-2.6, 0.5) | 0.7 | $T_{min}$ (°C) | 0.0 | -2.8<br>(-7.4, -1.8) | 2.1 |
| $T_{max}$ (°C) | 45.0 | 42.4<br>(38.4, 46.4) | 1.8 | $T_{max}$ (°C) | 45.0 | 40.3<br>(38.7, 41.9) | 0.7 |
| $q_0$ | 1.0000 | 0.0244<br>(0.0167, 0.0321) | 0.0035 | $A$ | 1.00 | 1.91<br>(1.84, 1.99) | 0.04 |
| | | | | $m$ | 1.00 | 0.33<br>(0.17, 0.48) | 0.07 |
| Step size (h) |  |  |  | 0.1 |  |  |  |
| Model evaluation |  |  |  |  |  |  |  |
| RSS | 0.0071 |  |  |  | 0.0155 |  |  |
| MSE | 0.0006 |  |  |  | 0.0016 |  |  |
| RMSE | 0.0253 |  |  |  | 0.0394 |  |  |
| AIC | -46.0602 |  |  |  | -29.8674 |  |  |
| AICc | -48.8602 |  |  |  | -29.8674 |  |  |
| BIC | -39.8276 |  |  |  | -22.7444 |  |  |

\* Int.: Initial guess; Est.: Estimation; SE: Standard error.
\*\* 95% CI: lower and upper 95% confidence intervals.

**Fig. 4.**
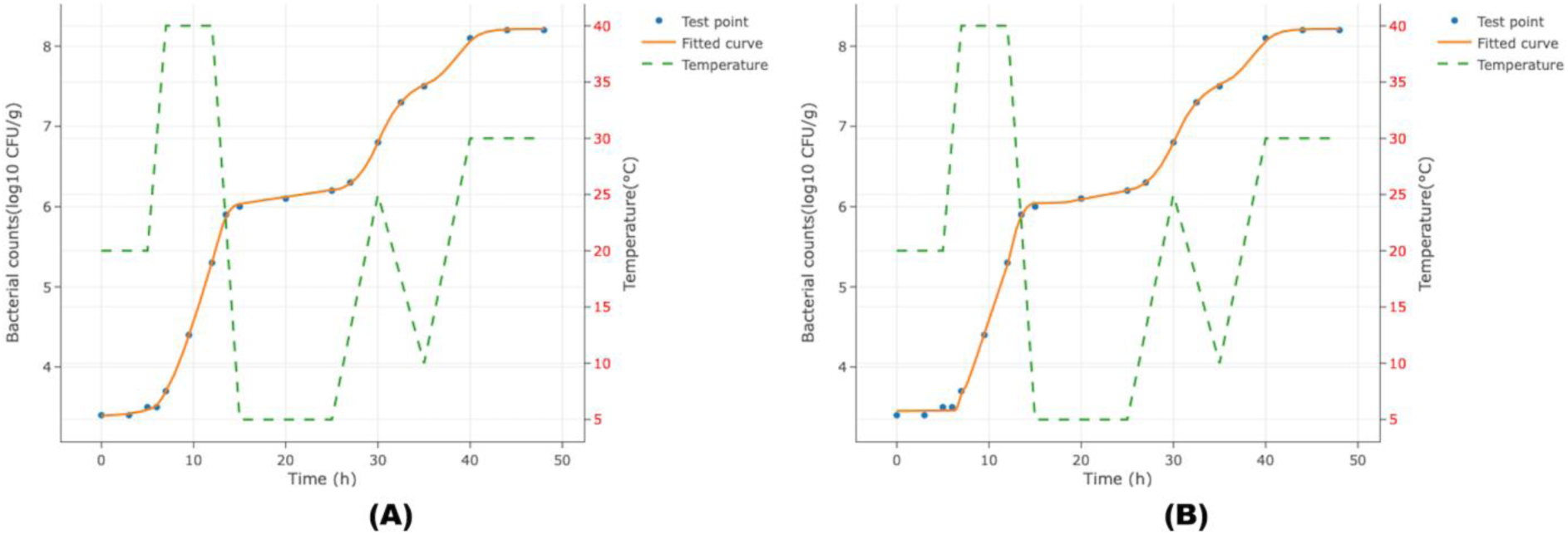
The fitted curve of Case IV with (A) the Baranyi-Cardinal parameter model and (B) the Huang-Cardinal parameter model downloaded from Microrisk Lab. (The blue dot represents the observed bacterial count, and the origin line represents the fitted curve.)

Similarly, with the microbial enumeration data and time-temperature profile in Case V, the non-isothermal inactivation fitting could be performed in Microrisk Lab (Fig. 5). Initial guesses of the estimable parameters were quoted from the primary study and listed in Tab.8, where the referenced temperature was fixed to 120°C (Garre, Fernández, Lindqvist, & Egea, 2017). As illustrated in Tab.8, the obtained estimations of Microrisk Lab are close to that of Bioinactivation FE. It should be noted, however, that numerical methods for the ordinary differential equations were different in these two tools. The **LSODA** solver in R package *‘deSolve’* (Soetaert, Petzoldt & Setzer, 2010) was introduced in Bioinactivation series to conduct the predictor-corrector method or backward differentiation formulae method for the dynamic model. In contrast, the Runge-Kutta method was provided by Microrisk Lab. These numerical methods have their own advantages and disadvantages respectively, but the choice might cause different truncation errors in a regression (Butcher, 2016). Thus, it is recommended to take care when using the evaluation indicators of AIC, AICc, and BIC provided from different modeling platforms for model comparison.

**Table 8.**
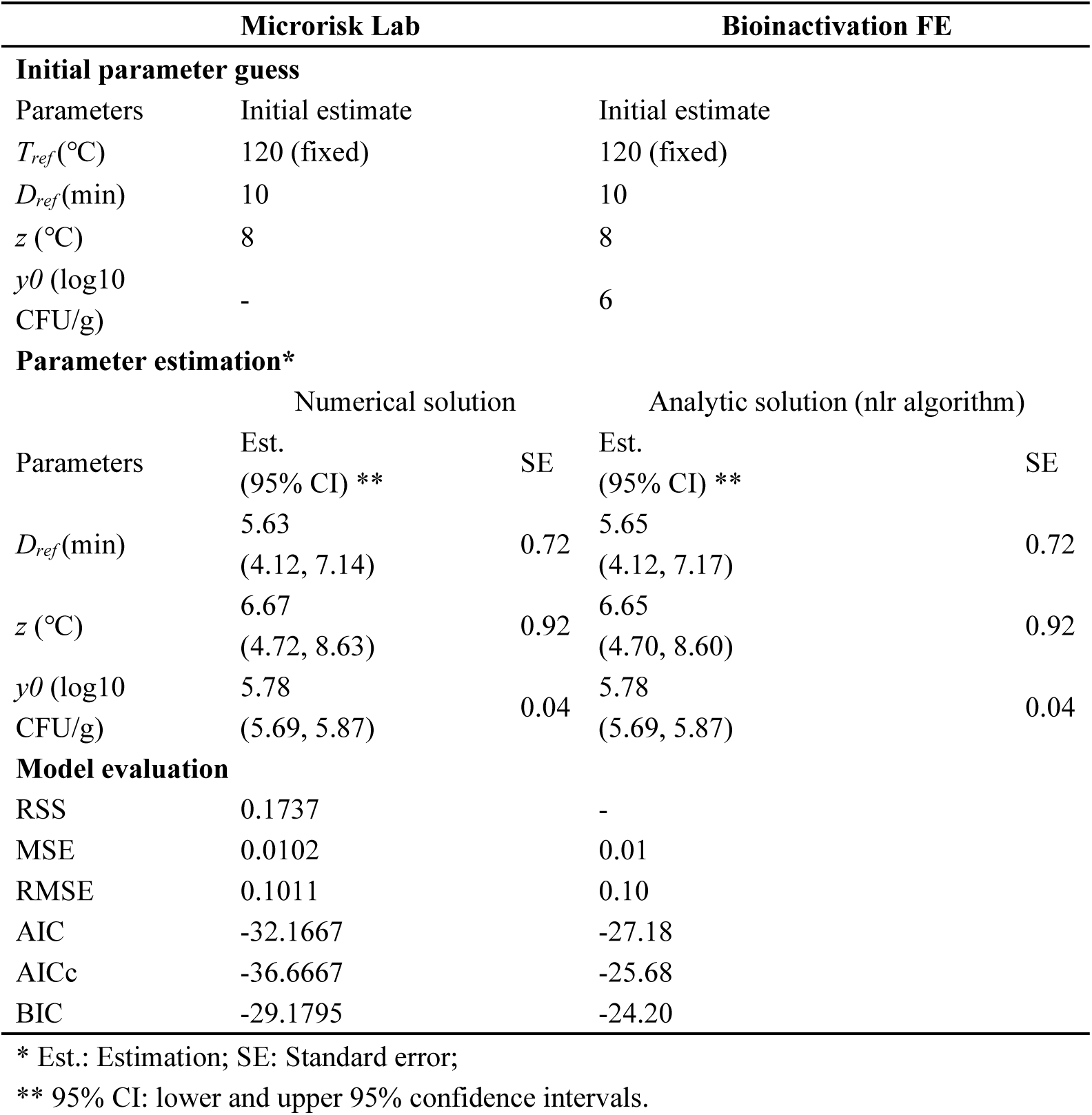
Comparison on non-isothermal inactivation fitting results of Microrisk Lab and Bioinactivation FE (Dynamic Bigelow model)

**Fig. 5.**
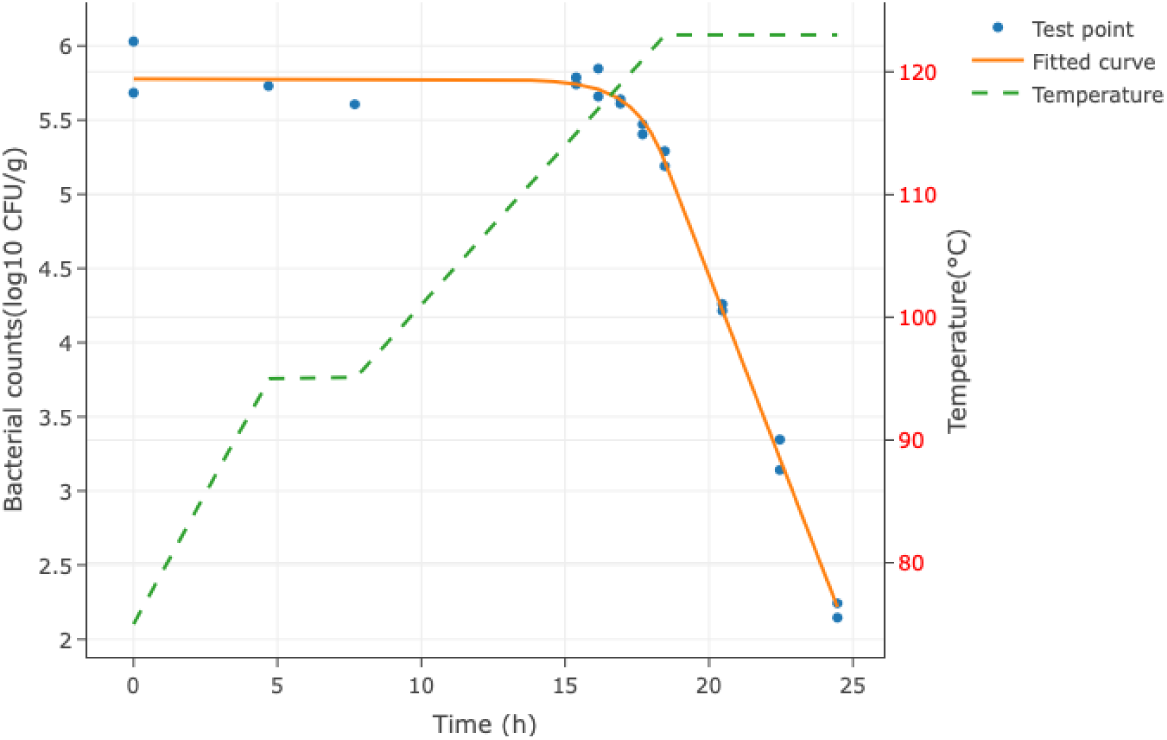
The fitted curve of Case V with the Dynamic Bigelow model downloaded from Microrisk Lab. (The blue dot represents the observed bacterial count, and the origin line represents the fitted curve.)

### 3.3. Comparison of the stochastic growth simulation

The stochastic type model is possible to be applied to the static simulation in Microrisk Lab by defining the distribution of different model variables. As previously mentioned, the behavior of microorganisms may be quite different when the population size decreases to the single-cell level. It is thus necessary to consider the uncertainty and variability of the cells during the simulation. In the referenced study of Case VI, Koutsoumanis & Lianou (2013) described the growth of the *S*. Typhimurium at the different single-cell level by establishing a stochastic model. Depending on the condition for the software of @Risk for Excel, the parameter setting of Microrisk Lab was listed in Tab.9, and the simulated results are presented in Fig.6(A). The probability distribution of the specific growth rate and the final bacterial concentration is provided with the mean value and standard deviation in Fig 6(C). According to the definition of the coefficient of variation (%CV = 100×standard deviation/mean) in the original study, the estimated %CV for *S*. Typhimurium final concentration is also around 25.5% in Microrisk Lab. The above result demonstrates that Microrisk Lab can perform a Monte-Carlo simulation for bacterial stochastic modeling. Moreover, Fig 6(D) shows the tornado graph of the sensitivity analysis on bacterial counts obtained by different associated parameters. Thus, restricted by the above settings, the uncertainty of the duration of growth time has a relatively higher impact (than other variables) on the bacterial count during the stochastic growth of *S*. Typhimurium single cell.

**Table 9.**
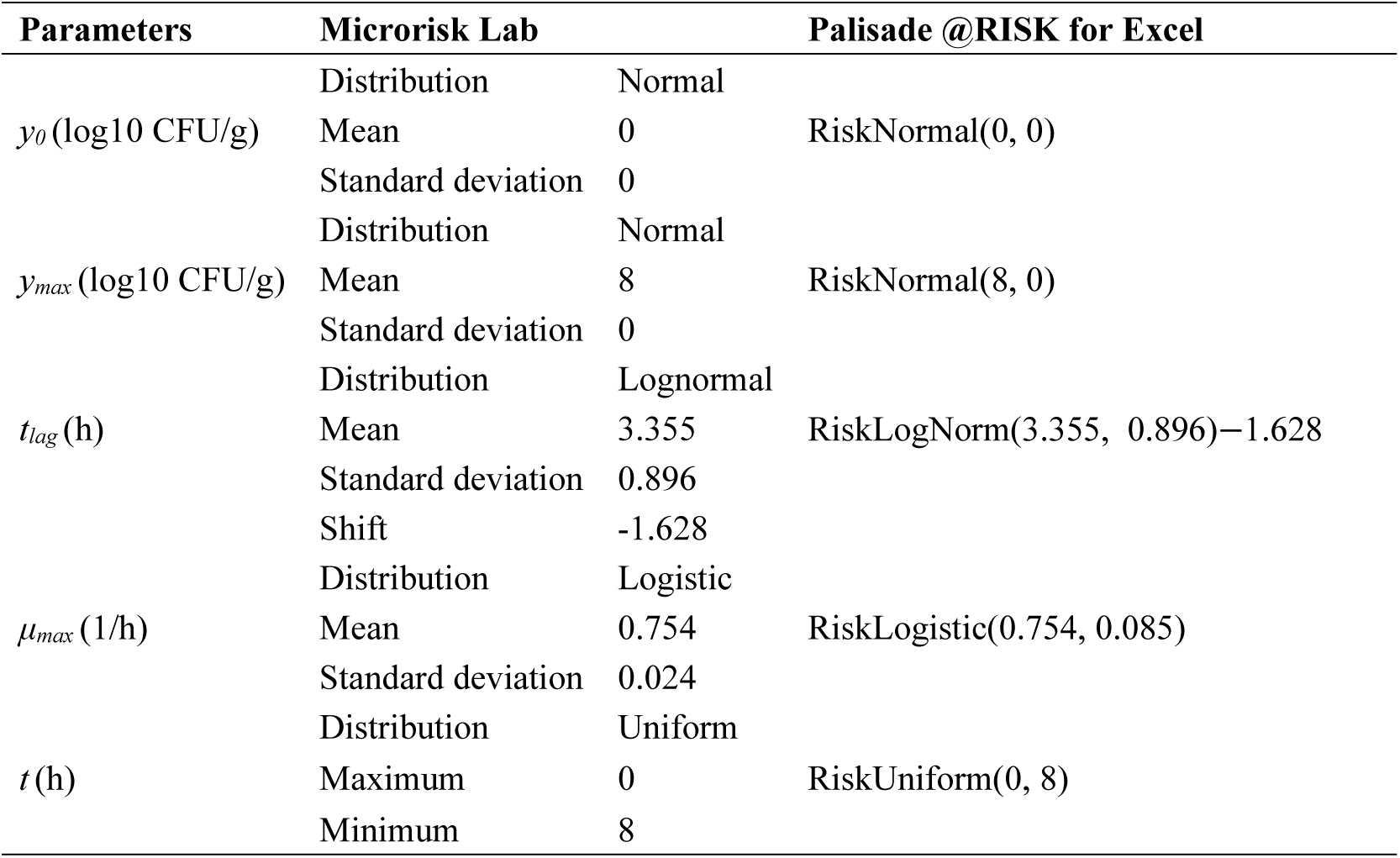
Stochastic growth simulation settings for Microrisk Lab and Palisade @RISK (Buchanan model)

**Fig. 6.**
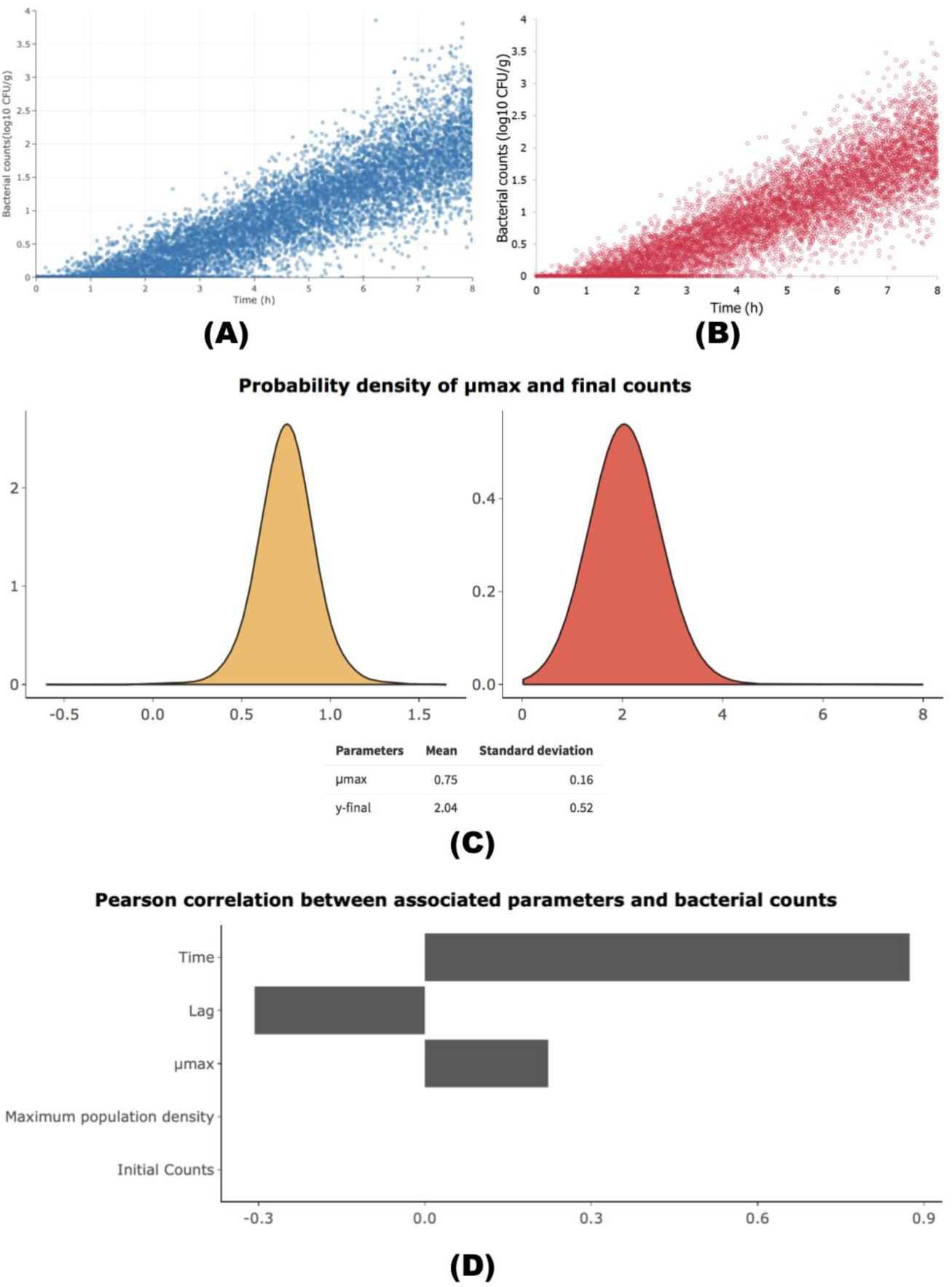
Monte-Carlo simulation of 1 cell growth with 10, 000 iterations in (A) Microrisk Lab, and (B) @RISK for Excel (adapted from Fig.7 of Koutsoumanis, & Lianou, 2013). (C) Simulated distribution of the maximum specific growth rate and final bacterial count. (D) Tornado graph of the sensitivity analysis between model variables and bacterial counts.

From the above cases, Microrisk Lab can be easily applied in microbial predictive modeling, however, functionalities should be improved to handle more practical modeling tasks. The model applicability could be expanded, for example, paying more attention to the impact of the interaction between different intrinsic or extrinsic factors on the microorganism. Algorithms involved in regression and simulation are also deserved to be developed for more options. Bioinactivation FE provides a good example for containing different fitting algorithms, while the functionality of fixed parameter could help users decide the estimable parameter (Garre et al, 2018). Meanwhile, Latin Hypercube sampling is a widely used method for the Monte-Carlo simulation in qualitative microbiological risk assessments (Ding et al., 2013; Membré & Boué, 2017; Dogan, Clarke, Mattos & Wang, 2019), which should be considered in our future update to improve the sampling efficiency.

## 4. Conclusions

In this study, a web-based freeware, Mircrorisk Lab, was introduced and used to validate its performance limited regression and simulation analysis in predictive microbiology. The interactive interface and simple manipulation logic help users readily obtain the modeling results. Practical examples elucidated that, in most cases, there was no statistical difference between the results obtained from Microrisk Lab and other existing modeling systems (except the online DMFit) in both regression and simulation studies. The new tool could provide more statistical results for the estimated parameter or evaluated indicator. Besides, it was also easy to perform the growth kinetic analysis under non-isothermal conditions without any coding skill in Microrisk Lab. This freeware might serve as a useful modeling tool and relevant educational resource for predictive modeling in microbiology.

## List of symbols

*Y*(*t*), *Y*_0_, *Y*_*max*_: the natural logarithm of real-time, initial, and maximum bacterial counts (ln CFU/g).
*y*(*t*), *y*_0_, *y*_*max*_: the 10-base logarithm of real-time, initial, and maximum bacterial counts (log10 CFU/g).
*y*_*res*_: the 10-base logarithm of the residual bacterial counts (log10 CFU/g).
*μ*_*max*_, *μ*_*opt*_: the maximum and optimal specific growth rate.
*k*_*max*_: the maximum specific inactivation rate.
*D*: the time of decimal reduction in inactivation.
*D*_*ref*_: the referenced decimal reduction time at *T*_*ref*_.
*t*_*lag*_: the time of lag in growth.
*S*_*l*_: the time of shoulder (or before inactivation) in inactivation.
*t*: the time point.
*t*_*max*_: the time when entering the stationary phase in growth.
*S*_*t*_: the time when entering the stationary phase in inactivation.
*T, pH, aw*: The temperature (°C), pH, and water activity at *t*.
*T*_*min*_, *T*_*opt*_, *T*_*max*_: the minimum, optimal, and maximum growth temperature (°C).
*T*_*ref*_: the referenced inactivation temperature (°C).
*pH*_*min*_, *pH*_*opt*_, *pH*_*max*_: the minimum, optimal, and maximum growth pH.
*aw*_*min*_, *aw*_*opt*_, *aw*_*max*_: the minimum, optimal, and maximum growth water activity.
*q*_0_: the initial physiological state of the inoculum in the Baranyi model.
*δ, p*: the coefficients in the Weibull model.
*δ*_*ref*_: the referenced *δ* value at *T*_*ref*_.
*a, b*: the coefficients in the square-root model.
*A, m*: the coefficients in the dynamic Huang model.
*z*: the coefficients of the bacterial thermal resistance (°C).

## Supplementary data

Supplementary data to this article is available.

## Acknowledgments

This work was supported by the National Key Research and Development Program of China (Grant No. 2019YFE0103800). We would like to thank Dr. Lihan Huang in the Eastern Regional Research Center of USDA, USA, for his helpful guidance on the R programming.

## Reference

Akaike, H. (1974). A new look at the statistical model identification. IEEE Transactions on Automatic Control, 19(6), 716–723. http://doi.org/10.1109/TAC.1974.1100705

Albert, I., & Mafart, P. (2005). A modified Weibull model for bacterial inactivation. International Journal of Food Microbiology, 100(1-3), 197–211. http://doi.org/10.1016/j.ijfoodmicro.2004.10.016

Alonso, A. A., Molina, I., & Theodoropoulos, C. (2014). Modeling bacterial population growth from stochastic single-cell dynamics. Applied and Environmental Microbiology, 80(17), 5241–5253. http://doi.org/10.1128/AEM.01423-14

Augustin, J.-C. (2011). Challenges in risk assessment and predictive microbiology of foodborne spore-forming bacteria. Food Microbiology, 28(2), 209–213. http://doi.org/10.1016/j.fm.2010.05.003

Augustin, J.-C., Ferrier, R., Hezard, B., Lintz, A., & Stahl, V. (2015). Comparison of individual-based modeling and population approaches for prediction of foodborne pathogens growth. Food Microbiology, 45, 205–215. http://doi.org/10.1016/j.fm.2014.04.006

Baranyi, J., & Buss da Silva, N. (2017). The use of predictive models to optimize risk of decisions. International Journal of Food Microbiology, 240, 19–23. http://doi.org/10.1016/j.ijfoodmicro.2016.10.016

Baranyi, J., & Roberts, T. A. (1995). Mathematics of predictive food microbiology. International Journal of Food Microbiology, 26(2), 199–218.

Baranyi, J., George, S. M., & Kutalik, Z. (2009). Parameter estimation for the distribution of single cell lag times. Journal of Theoretical Biology, 259(1), 24–30. http://doi.org/10.1016/j.jtbi.2009.03.023

Baty, F., & Delignette-Muller, M.-L. (2015). *nlsMicrobio*: Nonlinear regression in predictive microbiology. R package version 0.0-1. Available at: www.r-project.org.

Bigelow, W. D. (1921). The logarithmic nature of thermal death time curves. Journal of Infectious Diseases, 29(5), 528–536. http://doi.org/10.1093/infdis/29.5.528

Buchanan, R. L., & Golden, M. H. (1995). Model for the non-thermal inactivation of *Listeria monocytogenes* in a reduced oxygen environment. Food Microbiology, 12, 203–212. http://doi.org/10.1016/S0740-0020(95)80099-9

Buchanan, R. L., & Phillips, J. G. (1990). Response surface model for predicting the effects of temperature pH, sodium chloride content, sodium nitrite concentration and atmosphere on the growth of *Listeria monocytogenes*. Journal of Food Protection, 53(5), 370–376. http://doi.org/10.4315/0362-028x-53.5.370

Buchanan, R. L., Whiting, R. C., & Damert, W. C. (1997). When is simple good enough: a comparison of the Gompertz, Baranyi, and three-phase linear models for fitting bacterial growth curves. Food Microbiology, 14(4), 313–326. http://doi.org/10.1006/fmic.1997.0125

Burnham, K. P., & Anderson, D. R. (2003). Model Selection and Multimodel Inference. Springer Science & Business Media.

Busschaert, P., Geeraerd, A. H., Uyttendaele, M., & Van Impe, J. F. (2011). Sensitivity analysis of a two-dimensional quantitative microbiological risk assessment: Keeping variability and uncertainty separated. Risk Analysis, 31(8), 1295–1307. http://doi.org/10.1111/j.1539-6924.2011.01592.x

Butcher, J. C. (2016). Numerical methods for ordinary differential equations (Third edition). New York: Wiley.

Cattani, F., Dolan, K. D., Oliveira, S. D., Mishra, D. K., Ferreira, C. A. S., Periago, P. M., et al. (2016). One-step global parameter estimation of kinetic inactivation parameters for *Bacillus sporothermodurans* spores under static and dynamic thermal processes. Food Research International, 89(1), 614–619. http://doi.org/10.1016/j.foodres.2016.08.027

Chang, W., & Borges Ribeiro, B. (2019). *shinydashboard*: create dashboards with’Shiny’. R package version 0.7.1. Available at: www.r-project.org.

Cornu, M., Billoir, E., Bergis, H., Beaufort, A., & Zuliani, V. (2011). Modeling microbial competition in food: Application to the behavior of *Listeria monocytogenes* and lactic acid flora in pork meat products. Food Microbiology, 28(4), 639–647. http://doi.org/10.1016/j.fm.2010.08.007

Ding, T., Iwahori, J., Kasuga, F., Wang, J., Forghani, F., Park, M.-S., & Oh, D.-H. (2013). Risk assessment for *Listeria monocytogenes* on lettuce from farm to table in Korea. Food Control, 30(1), 190–199. http://doi.org/10.1016/j.foodcont.2012.07.014

Dogan, O. B., Clarke, J., Mattos, F., & Wang, B. (2019). A quantitative microbial risk assessment model of *Campylobacter* in broiler chickens: Evaluating processing interventions. Food Control, 100, 97–110. http://doi.org/10.1016/j.foodcont.2019.01.003

Dolan, K., Habtegebriel, H., Valdramidis V.P., & Mishra, D. (2015). Thermal processing and kinetic modeling of inactivation. In: Bakalis, S., Knoerzer, K., & Fryer, P.J. (Eds.), Modeling Food Processing Operations (pp. 37–66). Woodhead Publishing.

Draper, N. R., & Smith, H. (1998). Applied regression analysis. John Wiley & Sons.

Garre, A., Clemente-Carazo, M., Fernández, P. S., Lindqvist, R., & Egea, J. A. (2018). Bioinactivation FE: A free web application for modelling static and dynamic microbial inactivation. Food Research International, 112, 353–360. http://doi.org/10.1016/j.foodres.2018.06.057

Garre, A., Fernández, P. S., Lindqvist, R., & Egea, J. A. (2017). Bioinactivation: Software for modelling dynamic microbial inactivation. Food Research International, 93, 66–74. http://doi.org/10.1016/j.foodres.2017.01.012

Geeraerd, A. H., Herremans, C. H., & Van Impe, J. F. (2000). Structural model requirements to describe microbial inactivation during a mild heat treatment. International Journal of Food Microbiology, 59(3), 185–209.

Geeraerd, A. H., Valdramidis, V. P., & Van Impe, J. F. (2005). GInaFiT, a freeware tool to assess non-log-linear microbial survivor curves. International Journal of Food Microbiology, 102(1), 95–105. http://doi.org/10.1016/j.ijfoodmicro.2004.11.038

Geeraerd, A. H., Valdramidis, V. P., & Van Impe, J. F. (2006). Erratum to “GInaFiT, a freeware tool to assess non-log-linear microbial survivor curves” [Int. J. Food Microbiol. 102 (2005) 95–105]. International Journal of Food Microbiology, 110(3), 297–1. http://doi.org/10.1016/j.ijfoodmicro.2006.04.002

González, S. C., Possas, A., Carrasco, E., Valero, A., Bolívar, A., Posada-Izquierdo, G. D., et al. (2018). “MicroHibro”: A software tool for predictive microbiology and microbial risk assessment in foods. International Journal of Food Microbiology, 290, 226–236. http://doi.org/10.1016/j.ijfoodmicro.2018.10.007

Göransson, M., Nilsson, F., & Jevinger, Å. (2018). Temperature performance and food shelf - life accuracy in cold food supply chains–Insights from multiple field studies. Food Control, 86, 332–341. http://doi.org/10.1016/j.foodcont.2017.10.029

Hamner, B., Frasco, M., & LeDell, E. (2018). *Metrics*: Evaluation metrics for machine learning. R package version 0.1.4. Available at: www.r-project.org.

Huang, L. (2008). Growth kinetics of *Listeria monocytogenes* in broth and beef Frankfurters—Determination of lag phase duration and exponential growth rate under static conditions. Journal of Food Science, 73(5), E235–E242. http://doi.org/10.1111/j.1750-3841.2008.00785.x

Huang, L. (2014). IPMP 2013–a comprehensive data analysis tool for predictive microbiology. International Journal of Food Microbiology, 171, 100–107. http://doi.org/10.1016/j.ijfoodmicro.2013.11.019

Huang, L. (2016). Simulation and evaluation of different statistical functions for describing lag time distributions of a bacterial growth curve. Microbial Risk Analysis, 1, 47–55. http://doi.org/10.1016/j.mran.2015.08.002

Huang, L. (2017a). Dynamic identification of growth and survival kinetic parameters of microorganisms in foods. Current Opinion in Food Science, 14, 85–92. http://doi.org/10.1016/j.cofs.2017.01.013

Huang, L. (2017b). IPMP Global Fit–A one-step direct data analysis tool for predictive microbiology. International Journal of Food Microbiology, 262, 38–48. http://doi.org/10.1016/j.ijfoodmicro.2017.09.010

Huang, L., & Hwang, C.-A. (2017). Dynamic analysis of growth of *Salmonella Enteritidis* in liquid egg whites. Food Control, 80, 125–130. http://doi.org/10.1016/j.foodcont.2017.04.044

Huang, L., Hwang, C.-A., & Phillips, J. (2011). Evaluating the effect of temperature on microbial growth rate-The Ratkowsky and a Bělehrádek-type models. Journal of Food Science, 76(8), M547–M557. http://doi.org/10.1111/j.1750-3841.2011.02345.x

Hwang, C.-A., & Huang, L. (2018). Dynamic analysis of competitive growth of *Escherichia coli* O157:H7 in raw ground beef. Food Control, 93, 251–259. http://doi.org/10.1016/j.foodcont.2018.06.017

Iannetti, L., Salini, R., Sperandii, A. F., Santarelli, G. A., Neri, D., Di Marzio, V., et al. (2017). Predicting the kinetics of *Listeria monocytogenes* and *Yersinia enterocolitica* under dynamic growth/death-inducing conditions, in Italian style fresh sausage. International Journal of Food Microbiology, 240, 108–114. http://doi.org/10.1016/j.ijfoodmicro.2016.04.026

ICMSF, International Commission on Microbiological Specifications for Foods (1996). *Listeria monocytogenes*. In T. A. Robert, A. C. Baird-Parker, & R. B. Tompkin (Eds.), Microorganisms in foods 5: characteristics of microbial pathogens (pp. 148). London: Blackie Academic and Professional.

Koutsoumanis, K. P., & Lianou, A. (2013). Stochasticity in colonial growth dynamics of individual bacterial cells. Applied and Environmental Microbiology, 79(7), 2294–2301. http://doi.org/10.1128/AEM.03629-12

Koutsoumanis, K. P., Lianou, A., & Gougouli, M. (2016). Last developments in foodborne pathogens modeling. Current Opinion in Food Science, 8, 89–98. http://doi.org/10.1016/j.cofs.2016.04.006

Li, M., Huang, L., & Yuan, Q. (2017). Growth and survival of *Salmonella* Paratyphi A in roasted marinated chicken during refrigerated storage: Effect of temperature abuse and computer simulation for cold chain management. Food Control, 74, 17–24. http://doi.org/10.1016/j.foodcont.2016.11.023

Mafart, P., Couvert, O., Gaillard, S., & Leguérinel, I. (2002). On calcu lating sterility in thermal preservation methods: application of the Weibull frequency distribution model. International Journal of Food Microbiology, 72(1-2), 107–113.

Magalhães, R., Mena, C., Ferreira, V., Silva, J., Almeida, G., Gibbs, P., & Teixeira, P. (2014). *Listeria monocytogenes*. In: Y. Motarjemi, G. Moy, & E. Todd (Eds), Encyclopedia of Food Safety (pp. 450–461). Oxford: Elsevier’s Science & Technology.

Membré, J.-M., & Boué, G. (2018). Quantitative microbiological risk assessment in food industry: Theory and practical application. Food Research International, 106, 1132–1139. http://doi.org/10.1016/j.foodres.2017.11.025

Ndraha, N., Hsiao, H.-I., Vlajic, J., Yang, M.-F., & Lin, H.-T. V. (2018). Time-temperature abuse in the food cold chain: Review of issues, challenges, and recommendations. Food Control, 89, 12–21. http://doi.org/10.1016/j.foodcont.2018.01.027

Oscar, T. P. (2002). Development and validation of a tertiary simulation model for predicting the potential growth of *Salmonella* typhimurium on cooked chicken. International Journal of Food Microbiology, 76(3), 177–190. http://doi.org/10.1016/s0168-1605(02)00025-9

Peleg, M., & Corradini, M. G. (2011). Microbial growth curves: What the models tell us and what they cannot. Critical Reviews in Food Science and Nutrition, 51(10), 917–945. http://doi.org/10.1080/10408398.2011.570463

Plaza-Rodríguez, C., Thoens, C., Falenski, A., Weiser, A. A., Appel, B., Kaesbohrer, A., & Filter, M. (2015). A strategy to establish food safety model repositories. International Journal of Food Microbiology, 204, 81–90. http://doi.org/10.1016/j.ijfoodmicro.2015.03.010

Pouillot, R., Delignette-Muller, M., & Denis, J. (2017). *mc2d*: tools for two-dimensional Monte-Carlo simulations. R package version 0.1-18. Available at: www.r-project.org.

Press, W. H., Teukolsky, S. A., Vetterling, W. T., & Flannery, B. P. (2007). Chapter 17: Integration of ordinary differential equations. In: Numerical Recipes-the Art of Scientific Computing (pp.899–954). Cambridge: Cambridge University Press.

Presser, K. A., Ratkowsky, D. A., & Ross, T. (1997). Modelling the growth rate of *Escherichia coli* as a function of pH and lactic acid concentration. Applied and Environmental Microbiology, 63(6), 2355–2360.

Psomas, A. N., Nychas, G.-J., Haroutounian, S. A., & Skandamis, P. N. (2011). Development and validation of a tertiary simulation model for predicting the growth of the food microorganisms under dynamic and static temperature conditions. Computers and Electronics in Agriculture, 76(1), 119–129. http://doi.org/10.1016/j.compag.2011.01.013

Ratkowsky, D.A. (2003). Model fitting and uncertainty. In: R. McKellar, & X. Lu (Eds.), Modeling Microbial Responses in Foods (pp. 151–196). Boca Raton: CRC Press.

Ratkowsky, D. A., Lowry, R. K., McMeekin, T. A., Stokes, A. N., & Chandler, R. E. (1983). Model for bacterial culture growth rate throughout the entire biokinetic temperature range. Journal of Bacteriology, 154(3), 1222–1226.

Rawlings, J. O., Pantula, S. G., & Dickey, D. A. (2001). Chapter 7: Model development variable selection. In: Applied regression analysis: a research tool (pp.205-234). Springer Science & Business Media.

Ross, T., & McMeekin, T. A. (1994). Predictive microbiology. International Journal of Food Microbiology, 23(3-4), 241–264. http://doi.org/10.1016/0168-1605(94)90155-4

Rosso, L., & Robinson, T. P. (2001). A cardinal model to describe the effect of water activity on the growth of moulds. International Journal of Food Microbiology, 63(3), 265–273.

Rosso, L., Bajard, S., Flandrois, J. P., Lahellec, C., Fournaud, J., & Veit, P. (1996). Differential growth of *Listeria monocytogenes* at 4 and 8°C: Consequences for the shelf life of chilled products. Journal of Food Protection, 59(9), 944–949. http://doi.org/10.4315/0362-028X-59.9.944

Rosso, L., Lobry, J. R., & Flandrois, J. P. (1993). An unexpected correlation between cardinal temperatures of microbial growth highlighted by a new model. Journal of Theoretical Biology, 162(4), 447–463. http://doi.org/10.1006/jtbi.1993.1099

Rosso, L., Lobry, J. R., Bajard, S., & Flandrois, J. P. (1995). Convenient model to describe the combined effects of temperature and pH on microbial growth. Applied and Environmental Microbiology, 61(2), 610–616.

Schwarz, G. (1978). Estimating the dimension of a model. The Annals of Statistics, 6(2), 461–464. http://doi.org/10.1214/aos/1176344136

Tenenhaus-Aziza, F., & Ellouze, M. (2015). Software for predictive microbiology and risk assessment: A description and comparison of tools presented at the ICPMF8 Software Fair. Food Microbiology, 45(PB), 290–299. http://doi.org/10.1016/j.fm.2014.06.026

USDA, U.S. Department of Agriculture. (2016). Pathogen Modeling Program. https://www.ars.usda.gov/northeast-area/wyndmoor-pa/eastern-regional-research-center/residue-chemistry-and-predictive-microbiology-research/docs/pathogenmodeling-program/ Accessed 01 May 2018.

USDA. (2017). IPMP Dynamic Prediction. https://www.ars.usda.gov/northeast-area/wyndmoor-pa/eastern-regional-research-center/docs/ipmp-dynamic-prediction/ Accessed 01 May 2018.

van Boekel, M. A. J. S. (2002). On the use of the Weibull model to describe thermal inactivation of microbial vegetative cells. International Journal of Food Microbiology, 74(1-2), 139–159.

van Boekel, M. A. J. S., & Zwietering, M. H. (2007). Experimental design, data processing and model fitting in predictive microbiology. In: S. Brul, S. van Gerwen, M. Zwietering, (Eds.). Modeling microorganisms in food (pp. 38). Woodhead Publishing.

Van Impe, J. F., Nicolaï, B. M., Martens, T., De Baerdemaeker, J., & Vandewalle, J. (1992). Dynamic mathematical model to predict microbial growth and inactivation during food processing. Applied and Environmental Microbiology, 58(9), 2901–2909.

Vimont, A., Vernozy-Rozand, C., Montet, M. P., Lazizzera, C., Bavai, C., & Delignette-Muller, M. L. (2006). Modeling and predicting the simultaneous growth of *Escherichia coli* O157:H7 and ground beef background microflora for various enrichment protocols. Applied and Environmental Microbiology, 72(1), 261–268. http://doi.org/10.1128/AEM.72.1.261-268.2006

Wang, X., Devlieghere, F., Geeraerd, A., & Uyttendaele, M. (2017). Thermal inactivation and sublethal injury kinetics of *Salmonella enterica* and *Listeria monocytogenes* in broth versus agar surface. International Journal of Food Microbiology, 243, 70–77. http://doi.org/10.1016/j.ijfoodmicro.2016.12.008

Whiting, R. C., & Buchanan, R. L. (1993). A classification of models in predictive microbiology. Food Microbiology, 10, 175–177.

WHO, World Health Organization (2015). WHO estimates of the global burden of foodborne diseases: foodborne disease burden epidemiology reference group 2007-2015. http://apps.who.int/iris/handle/10665/199350/ Accessed 01 May 2018.

Wickham, H., Chang, W., Henry, L., Pedersen, T.L., Takahashi, K., Wilke, C., & Woo, K. (2019). *ggplot2*: Create elegant data visualisations using the grammar of graphics. R package version 3.1.1. Available at: www.r-project.org.

Zwietering, M. H., Jongenburger, I., Rombouts, F. M., & van ‘t Riet, K. (1990). Modeling of the bacterial growth curve. Applied and Environmental Microbiology, 56(6), 1875–1881.

