## Supplementary material for "Microrisk Lab: an online freeware for predictive microbiology": S.1-User Manual of Microrisk Lab.pdf

#### User Manual

*Revised at September 2019*

*Version 1.0*

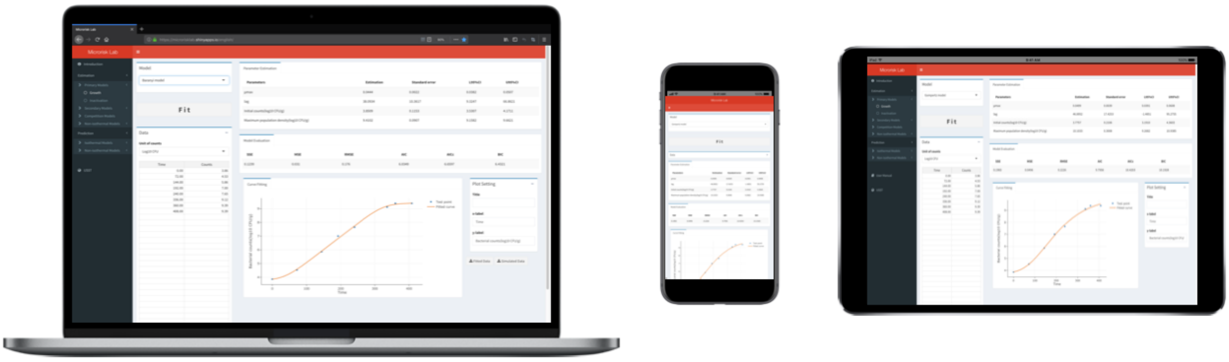

### Disclaimer and Support

#### Disclaimer

Microrisk Lab and this manual provide NO WARRANTY. This tool is free to use but only for research purposes. It is not permitted to include Microrisk Lab in any other application. We would very appreciate acknowledgement if the tool is used.

#### Feedback

If you have any suggestion or for technical questions for Microrisk Lab, please contact the developer and maintainer Yangtai Liu, Your comments are highly appreciated.

#### Document Revisions

| Date | Version | Document Changes |
|---|---|---|
| 01/09/2017 | Beta 0.1 | Initial draft |
| 01/01/2018 | 1.0 | Draft for updated version |
| 09/29/2019 | 1.0 | Revised draft |

### 1 List of symbols

| | |
|---|---|
| $Y(t)$, $Y_0$, $Y_{max}$ | the natural logarithm of real-time, initial, and maximum bacterial counts (ln CFU/g). |
| $y(t)$, $y_0$, $y_{max}$ | the 10-base logarithm of real-time, initial, and maximum bacterial counts (log10 CFU/g). |
| $y_{res}$ | the 10-base logarithm of the residual bacterial counts (log10 CFU/g). |
| $\mu_{max}$, $\mu_{opt}$ | the maximum and optimal specific growth rate. |
| $k_{max}$ | the maximum specific inactivation rate. |
| $D$ | the time of decimal reduction in inactivation. |
| $D_{ref}$ | the referenced decimal reduction time at $T_{ref}$. |
| $t_{lag}$ | the time of lag in growth. |
| $S_l$ | the time of shoulder (or before inactivation) in inactivation. |
| $t$ | the time point. |
| $t_{max}$ | the time when entering the stationary phase in growth. |
| $S_t$ | the time when entering the stationary phasein inactivation. |
| $T$, $pH$, $aw$ | The temperature (℃), pH, and water activity at $t$. |
| $T_{min}$, $T_{opt}$, $T_{max}$ | the minimum, optimal, and maximum growth temperature (℃). |
| $T_{ref}$ | the referenced inactivation temperature (℃). |
| $pH_{min}$, $pH_{opt}$, $pH_{max}$ | the minimum, optimal, and maximum growth pH. |
| $aw_{min}$, $aw_{opt}$, $aw_{max}$ | the minimum, optimal, and maximum growth water activity. |
| $q_0$ | the initial physiological state of the inoculum in the Baranyi model. |
| $\delta$, $p$ | the coefficients in the Weibull model. |
| $\delta_{ref}$ | the referenced $\delta$ value at $T_{ref}$. |
| $a$, $b$ | the coefficients in the square-root model. |
| $A$, $m$ | the coefficients in the dynamic Huang model. |
| $z$ | the coefficients of the bacterial thermal resistance (℃). |

### 2 Unit

The unit of the bacterial count and time related variables can be defined by the user. The unit of predicted counting outputs will be transferred into 10-base logarithm. Note that the unit of the specific (growth/ inactivation) rate is a natural logarithm combined with a unit of time, for example, ln CFU/g/h or ln CFU/g/min.

### 3 Programing basics

Microrisk Lab is developed by the open-source language R (version 3.5.1 for Mac OS X; http://ww.r-project.org). All users are free to access and use this tool through the browser of any internet-connected device by the following links:

http://microrisklab.shinyapps.io/english (in English)

http://microrisklab.shinyapps.io/chinese (in Chinese)

The operation of this Microrisk Lab must depend on certain developed R packages, which were listed in Tab.1. All the required packages have been hosted and deployed in the Shinyapps.io sever (https://www.shinyapps.io).

Tab.1 Imported R packages in Microrisk Lab

| Package name | Version | Reference | Purpose |
|---|---|---|---|
| *ggplot2* | 3.3.1 | Wickham et al. | to generate visualized plots for output |
| *mc2d* | 0.1-18 | Pouillot et al. | to generate certain distribution for output |
| *Metrics* | 0.1.4 | Hamner et al. | to calculate statistical indicators for output |
| *plotly* | 4.9.0 | Sievert et al. | to generate interactive plots for output |
| *rhandsontable* | 0.3.7 | Owen et al. | to build interactive table for input |
| *shiny* | 1.0.5 | Chang et al. | to establish and upload the shiny app |
| *shinyalert* | 1.0 | Attali et al. | to pop the error alert for input and output |
| *shinydashboard* | 0.7.1 | Chang et al. | to build the interactive interface |
| *shinyWidgets* | 0.4.8 | Perrier et al. | to build the interactive interface |
| *stats* | 3.4.3 | - | to realize the regression analysis |

Microrisk Lab can be also used on computers without internet connection when installed locally. In this case, please contact the developer.

### 4 Functionalities included in Microrisk Lab

Microrisk Lab includes the following functions:

- Kinetic analysis of microbial isothermal growth
- Kinetic analysis of microbial non-isothermal growth
- Kinetic analysis of microbial isothermal inactivation
- Kinetic analysis of microbial non-isothermal inactivation
- Kinetic analysis of two-flora isothermal competition growth
- Secondary modeling of specific growth rate vs. temperature, pH and Aw.
- Deterministic/ Stochastic simulation for microbial isothermal growth
- Deterministic simulation for microbial isothermal growth
- Deterministic/ Stochastic simulation for microbial isothermal inactivation
- Deterministic simulation for microbial isothermal inactivation
- Output interactive plots of the fitted and predicted curve.
- Output estimated results (estimates, standard error, and 95% confidential intervals) and multiple statistical indicators (RSS, MSE, RMSE, AIC, AICc, BIC, $R^2$, and Adjusted $R^2$) with respect to the experimental data in the 'Estimation' module.
- Output simulated bacterial counts or the distribution of the specific rate and final bacterial counts in the 'Simulation' module.
- Output correlation analysis between model parameters and simulated bacterial counts in the stochastic simulation.

### 5 Layout of Microrisk Lab

Fig.1 shows the page structure when loading in the Microrisk Lab via the browser in different devices. Users may switch the task by the main menu on the left side. In the setting panel, user can input the experimental data and choose the model here. The result panel will provide the estimated (predicted) values, statistical results, and interactive plots according to the setting.

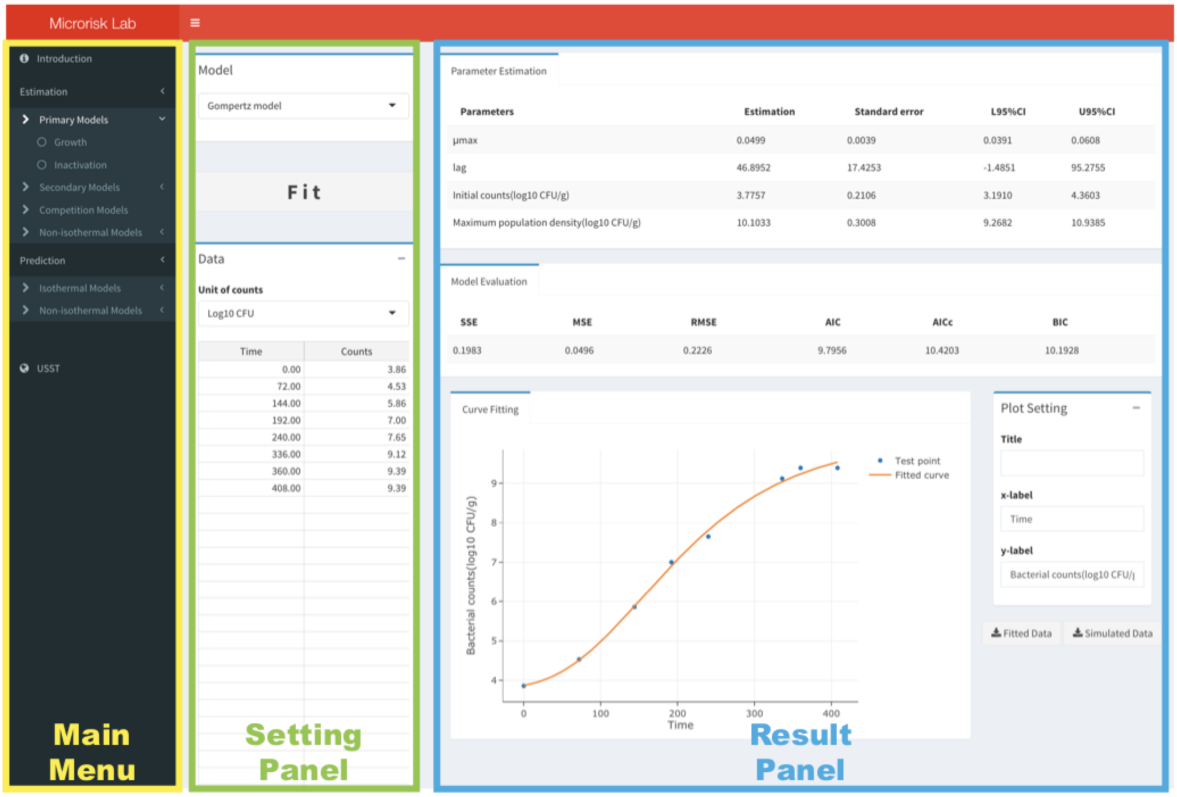

Fig.1 Typical layout of Microrisk Lab.

### 6 Estimation module of Microrisk Lab

The estimation module allows to solve multiple inverse problems in predictive microbiology, including ① isothermal growth fitting, ② isothermal inactivation fitting, ③④⑤ secondary model fitting, ⑥ two flora competition growth fitting, ⑦ non-isothermal growth fitting, and ⑧ non-isothermal inactivation fitting (Fig.2).

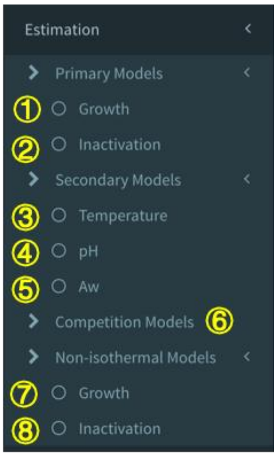

Fig.2 Different sections of model fitting in the estimation module.

#### Practical example Ⅰ - Isothermal growth fitting

(1) In this case, a group of *Listera monocytogenes*/ *innocua* growth in tryptose phosphate broth (TPB) obtained from the ComBase database (www.combase.cc, ComBase ID: LM127_11) was used as the test dataset for the growth fitting.

(2) Choose ① the 'Growth' in the section of the 'Primary Models', and the setting panel of isothermal growth model will show up (Fig.3).

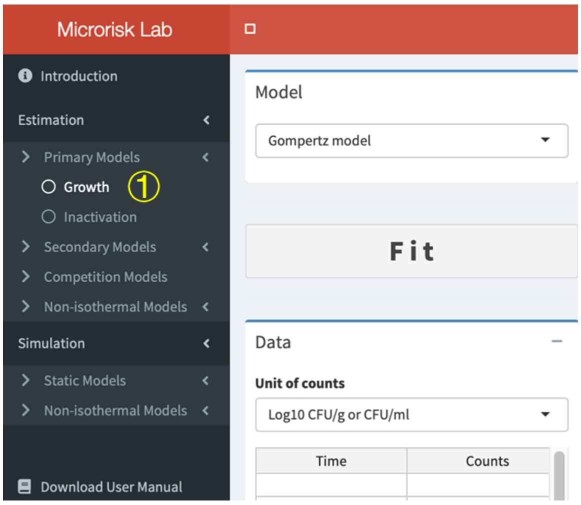

Fig.3 Layout of a section of the 'Estimation' module.

(3) The experimental data can be ① directly typed (or ② copied from other table files) in the ‘Data’ box. Specifically, ③ the unit of bacterial counts should be confirmed by the user. If the inputted observations are more than 30, please ④ right click the mouse or ⑤ drag the last column to add additional columns (Fig.4).

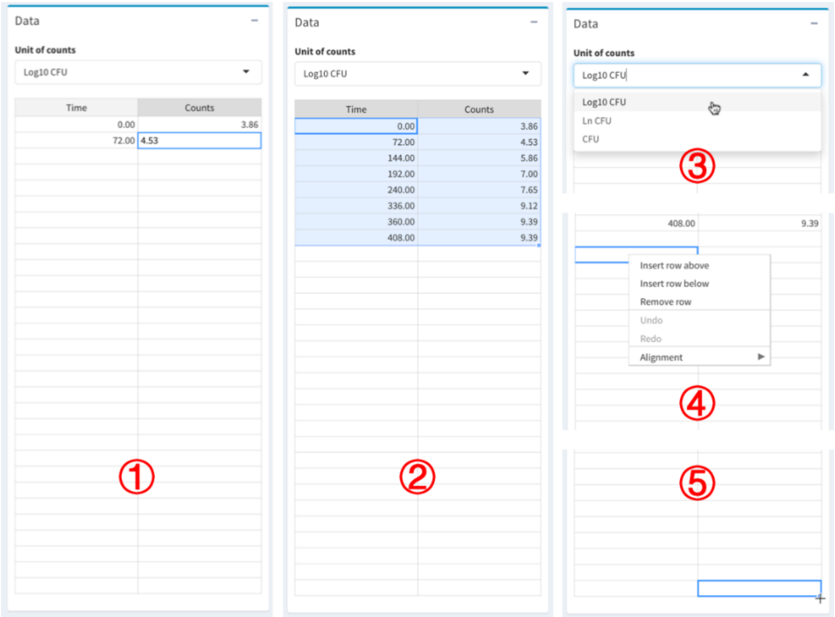

Fig.4 Boxes for the data input and unit selection.

(4) After entering the data for model fitting, the growth model can be selected in the ‘Model’ list (Fig.5).

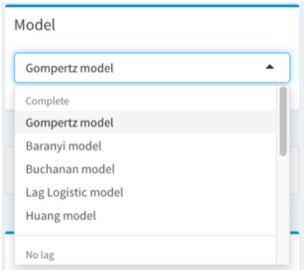

Fig.5 Box for the growth model selection

(5) Click ① the ‘Fitting’ button. After a necessary loading time, if the regression can be solved successfully, the ② estimated result, ③ evaluated result, and ④ interactive plot of the observation and fitted curve will show in the result panel (Fig.6A). Otherwise, a popup message will appear for the regression warning, which means that the non-linear regression is failed (Fig.6B). In this case, please re-check the unit of bacterial counts and the model selection.

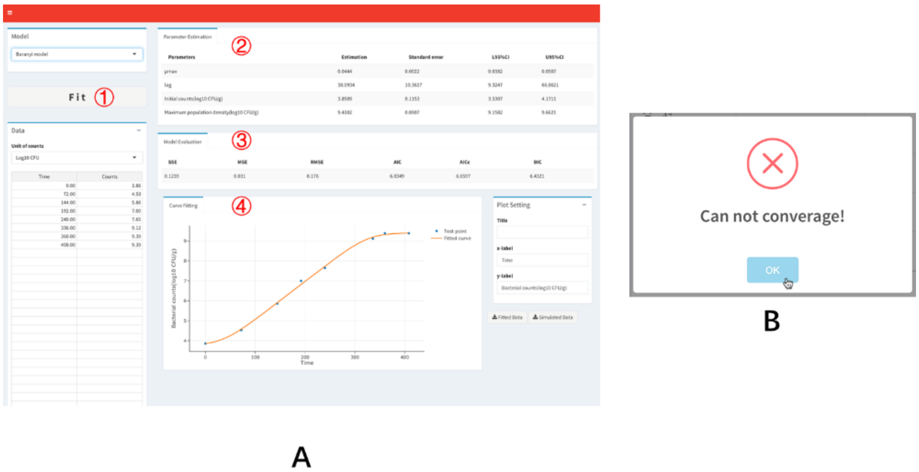

Fig.6 Layout of the interface after model fitting.

(6) The observed and predicted value can be viewed on ① the interactive plot. The observed data or fitting curve can be omitted from the plot by clicking ② the legend. Meanwhile, it is easy to edit the axis detail (③ the range and ④ title) of the interactive plot in real-time by ⑤ the box of 'Plot Setting'. After all, the plot is adjustable and downloadable by using ⑥ the 'Plotly toolbox'. Meanwhile, ⑦ the fitted and simulated data can be saved as the '.csv' file for comparison and further model development (Fig.7).

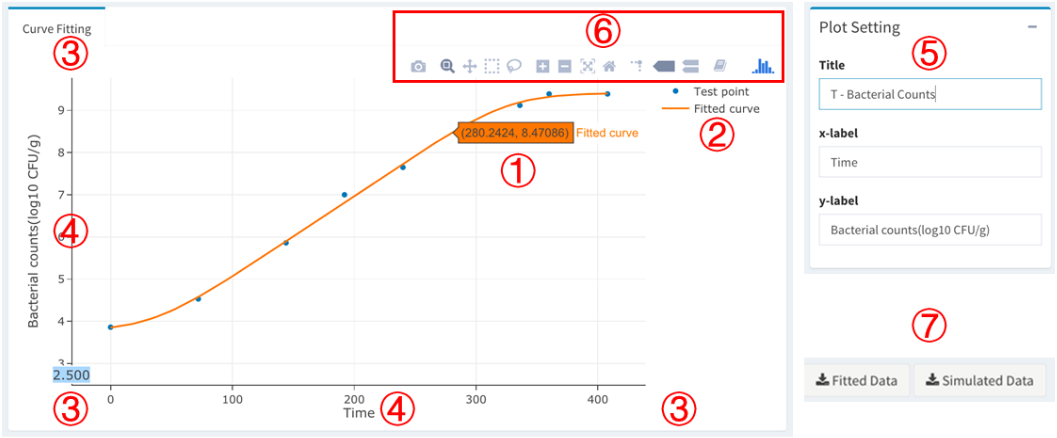

Fig.7 The interactive plot and the editorial box.

(7) For this case, the estimated result of parameters by using different growth models are listed in Tab.1, Tab.2, and Tab.3.

Tab.1 Static growth fitting results of the complete model in Microrisk Lab

| | Gompertz | | Baranyi | | Buchanan | | Lag Logistic | | Huang | |
|---|---|---|---|---|---|---|---|---|---|---|
| **Parameter estimation*** | | | | | | | | | | |
| Parameters | Est. (95% CI) ** | SE | Est. (95% CI) ** | SE | Est. (95% CI) ** | SE | Est. (95% CI) ** | SE | Est. (95% CI) ** | SE |
| $y_0$ (log10 CFU/g) | 3.78 (3.19, 4.36) | 0.21 | 3.85 (3.53, 4.17) | 0.12 | 3.86 (3.44, 4.28) | 0.15 | 3.86 (3.53, 4.17) | 0.11 | 3.86 (3.55, 4.17) | 0.11 |
| $y_{max}$ (log10 CFU/g) | 10.10 (9.27, 10.94) | 0.30 | 9.41 (9.16, 9.66) | 0.09 | 9.30 (9.06, 9.54) | 0.09 | 9.42 (9.16, 9.66) | 0.09 | 9.42 (9.18, 9.66) | 0.09 |
| $t_{lag}$ (h) | 38.09 (-1.49, 95.28) | 17.43 | 38.09 (9.32, 66.86) | 10.36 | 36.01 (3.06, 68.97) | 11.87 | 35.52 (9.32, 66.86) | 8.59 | 35.52 (11.69, 59.36) | 8.59 |
| $\mu_{max}$ (1/h) | 0.050 (0.04, 0.06) | 0.004 | 0.044 (0.038, 0.051) | 0.002 | 0.044 (0.036, 0.052) | 0.003 | 0.044 (0.038, 0.051) | 0.002 | 0.044 (0.039, 0.049) | 0.00 |
| **Model evaluation** | | | | | | | | | | |
| RSS | 0.1983 | | 0.1239 | | 0.0897 | | 0.1128 | | 0.1128 | |
| MSE | 0.0496 | | 0.0310 | | 0.0224 | | 0.0282 | | 0.0282 | |
| RMSE | 0.2226 | | 0.1760 | | 0.1497 | | 0.1679 | | 0.1679 | |
| AIC | 7.7956 | | 4.0349 | | -5.2237 | | 3.2840 | | 3.2840 | |
| AICc | 13.1289 | | 9.3683 | | 0.1096 | | 8.6173 | | 8.6173 | |
| BIC | 8.1133 | | 4.3527 | | -4.9059 | | 3.6018 | | 3.6018 | |

* Est.: Estimation; SE: Standard error.

** 95%CI: lower and upper 95% confidence intervals.

Tab.2 Static growth fitting results of the no lag model and linear model in Microrisk Lab

| | No lag Logistic | | No lag Buchanan | | Linear | |
|---|---|---|---|---|---|---|
| **Parameter estimation*** | | | | | | |
| Parameters | Est. (95% CI) ** | SE | Est. (95% CI) ** | SE | Est. (95% CI) ** | SE |
| $y_0$ (log10 CFU/g) | 3.60 (3.18, 4.02) | 0.16 | 3.65 (3.27, 4.03) | 0.15 | 3.81 (3.25, 4.37) | 0.23 |
| $y_{max}$ (log10 CFU/g) | 9.51 (8.98, 10.04) | 0.21 | 9.39 (9.02, 9.76) | 0.14 | - - | - |
| $\mu_{max}$ (1/h) | 0.039 (0.033, 0.045) | 0.002 | 0.038 (0.033, 0.042) | 0.002 | 0.035 (0.030, 0.040) | 0.002 |
| **Model evaluation** | | | | | | |
| $R^2$ | - | | - | | 0.9793 | |
| Adjusted $R^2$ | - | | - | | 0.9751 | |
| RSS | 0.5122 | | 0.2037 | | 0.3017 | |
| MSE | 0.1024 | | 0.0407 | | 0.0503 | |
| RMSE | 0.3201 | | 0.2019 | | 0.2242 | |
| AIC | 13.3871 | | -0.6606 | | 7.1519 | |
| AICc | 13.3871 | | -0.6606 | | 5.5519 | |
| BIC | 13.6254 | | -0.4222 | | 7.3108 | |

* Est.: Estimation; SE: Standard error.

** 95%CI: lower and upper 95% confidence intervals.

Tab.3 Static growth fitting results of the reduced model in Microrisk Lab

| | Reduced Baranyi | | Reduced Buchanan | | Reduced Huang | |
|---|---|---|---|---|---|---|
| **Parameter estimation*** | | | | | | |
| Parameters | Est. (95% CI) ** | SE | Est. (95% CI) ** | SE | Est. (95% CI) ** | SE |
| $y_0$ (log10 CFU/g) | 3.83 (2.90, 4.82) | 0.37 | 3.86 (3.53, 4.17) | | 3.86 (3.53, 4.17) | |
| $t_{lag}$ (h) | 2.13 (-78.88, 90.42) | 32.93 | 5.77 (9.32, 66.86) | 8.59 | 5.77 (9.32, 66.86) | 8.59 |
| $\mu_{max}$ (1/h) | 0.035 (0.028, 0.042) | 0.003 | 0.035 (0.038, 0.051) | | 0.035 (0.038, 0.051) | |
| **Model evaluation** | | | | | | |
| RSS | 1.5981 | | 0.6904 | | 1.5897 | |
| MSE | 0.3196 | | 0.1381 | | 0.3179 | |
| RMSE | 0.5653 | | 0.3716 | | 0.5639 | |
| AIC | 22.4900 | | 9.1037 | | 22.4482 | |
| AICc | 22.4900 | | 9.1037 | | 22.4482 | |
| BIC | 22.7283 | | 9.3420 | | 22.6865 | |

* Est.: Estimation; SE: Standard error.

** 95%CI: lower and upper 95% confidence intervals.

#### Practical example Ⅱ - Isothermal inactivation fitting

(1) In this case, a group of *Escherichia coli* inactivation in broth with 5% ethanol and 11200 ppm lactic acid (pH=3.8) at 5°C obtained from the ComBase database (ComBase ID: CA_Ec025) was used as the test dataset for the inactivation fitting.

(2) Choose the ② 'Inactivation' in the section of the 'Primary Models' (see Fig.2), and the setting panel of isothermal inactivation model will show up. The steps of importing data and obtaining results are similar to the isothermal growth fitting (see Fig.4 to Fig.7).

(3) For this case, the estimated result of parameters by using different growth model are listed in Tab.4, Tab.5 and Tab.6.

Tab.4 Static inactivation fitting results of the Geeraerd model in Microrisk Lab

| | **Complete Geeraerd** | | **No shoulder Geeraerd** | | **No tail Geeraerd** | |
|---|---|---|---|---|---|---|
| **Parameter estimation*** | | | | | | |
| Parameters | Est. (95% CI) ** | SE | Est. (95% CI) ** | SE | Est. (95% CI) ** | SE |
| $y_0$ (log10 CFU/g) | 8.97 (6.37, 11.56) | 0.20 | 9.10 (8.24, 9.97) | 0.20 | 9.02 (7.86, 10.18) | 0.27 |
| $y_{res}$ (log10 CFU/g) | 6.71 (3.42, 9.99) | 0.26 | 6.52 (4.34, 8.71) | 0.51 | - - | - |
| $S_l$ (h) | 1.65 (-11.56, 14.86) | 1.04 | | | -0.01 (-8.44, 8.41) | 1.96 |
| $k_{max}$ (1/h) | 1.184 (-3.944, 6.311) | 0.404 | 0.805 (0.099, 1.511) | 0.164 | 0.690 (-0.084, 1.464) | 0.180 |
| **Model evaluation** | | | | | | |
| RSS | 0.0429 | | 0.1047 | | 0.1480 | |
| MSE | 0.0429 | | 0.0523 | | 0.0740 | |
| RMSE | 0.2071 | | 0.2288 | | 0.2720 | |
| AIC | -1.6056 | | 0.8567 | | 2.5894 | |
| AICc | - | | 18.8567 | | 20.5894 | |
| BIC | -3.1678 | | -0.3150 | | 1.4177 | |

* Est.: Estimation; SE: Standard error.

** 95%CI: lower and upper 95% confidence intervals.

Tab.5 Static inactivation fitting results of the three/ two phase model in Microrisk Lab

| | **Three-phase** | | **No shoulder two-phase** | | **No tail two-phase** | |
|---|---|---|---|---|---|---|
| **Parameter estimation*** | | | | | | |
| Parameters | Est. (95% CI) ** | SE | Est. (95% CI) ** | SE | Est. (95% CI) ** | SE |
| $y_0$ (log10 CFU/g) | 9.00 (6.93, 11.07) | 0.16 | 9.11 (8.45, 9.77) | 0.15 | 9.00 (7.83, 10.17) | 0.27 |
| $y_{res}$ (log10 CFU/g) | 6.80 (4.73, 8.87) | 0.16 | 6.80 (6.01, 7.59) | 0.18 | - - | - |
| $S_l$ (h) | 0.92 (-7.32, 9.15) | 0.65 | - - | - | 0.16 (-5.78, 6.11) | 1.38 |
| $k_{max}$ (1/h) | 0.921 (-0.768, 2.610) | 0.133 | 0.794 (0.389, 1.200) | 0.094 | 0.702 (0.102, 1.303) | 0.140 |
| **Model evaluation** | | | | | | |
| RSS | 0.0267 | | 0.0670 | | 0.1470 | |
| MSE | 0.0267 | | 0.0335 | | 0.0735 | |
| RMSE | 0.1633 | | 0.1830 | | 0.2711 | |
| AIC | -3.9795 | | -1.3731 | | 2.5556 | |
| AICc | - | | 16.6269 | | 20.5556 | |
| BIC | -5.5418 | | -2.5448 | | 1.3839 | |

* Est.: Estimation; SE: Standard error.

** 95%CI: lower and upper 95% confidence intervals.

Tab.6 Static inactivation fitting results of the Weibull model and Bigelow model in Microrisk Lab

| | Webull-tail | | Webull | | Bigelow | |
|---|---|---|---|---|---|---|
| **Parameter estimation*** | | | | | | |
| Parameters | Est.<br>(95% CI) ** | SE | Est.<br>(95% CI) ** | SE | Est.<br>(95% CI) ** | SE |
| $y_0$ (log10 CFU/g) | 8.96<br>(6.94, 10.97) | 0.16 | 9.04<br>(7.89, 10.19) | 0.27 | 9.02<br>(8.47, 9.57) | 0.17 |
| $y_{res}$ (log10 CFU/g) | 8.96<br>(4.53, 8.98) | 0.18 | -<br>- | - | | |
| $p$ | 1.73<br>(-5.05, 8.51) | 0.53 | 3.21<br>(-1.58, 8.00) | 1.11 | | |
| $\delta$ | 3.63<br>(-2.65, 9.90) | 0.49 | 0.96<br>(-0.38, 2.30) | 0.31 | | |
| $D$ (h) | -<br>- | - | -<br>- | - | 3.33<br>(2.09, 4.58) | 0.39 |
| **Model evaluation** | | | | | | |
| $R^2$ | - | | - | | 0.9605 | |
| Adjusted $R^2$ | - | | - | | 0.9408 | |
| RSS | 0.0271 | | 0.1467 | | 0.1480 | |
| MSE | 0.0271 | | 0.0733 | | 0.0493 | |
| RMSE | 0.1648 | | 0.2708 | | 0.2221 | |
| AIC | -3.8901 | | 2.5449 | | 0.5895 | |
| AICc | - | | 20.5449 | | 2.5895 | |
| BIC | -5.4524 | | 1.3732 | | -0.1916 | |

* Est.: Estimation; SE: Standard error.

** 95%CI: lower and upper 95% confidence intervals.

#### Practical example Ⅲ - Temperature secondary model fitting

(1) In this case, a study on the maximum specific growth rate of *Salmonella* Typhimurium (ATCC 14028) in chicken breast (Oscar, 2002) was cited for fitting different secondary models.

(2) Choose the ③ 'Temperature' in the section of the 'Secondary Models' (see Fig.2), and the setting panel of temperature secondary model will show up. The steps of importing data and obtaining results are similar to the isothermal growth fitting (see Fig.4 to Fig.7).

(3) For this case, the estimated result of parameters by using different temperature secondary model are listed in Tab.7 and Tab.8.

Tab.7 Specific maximum growth rate fitting results of the suboptimal temperature secondary model in Microrisk Lab

| | Square-root model | | Huang square-root model | |
|---|---|---|---|---|
| **Parameter estimation*** | | | | |
| Parameters | Est.<br>(95% CI) ** | SE | Est.<br>(95% CI) ** | SE |
| $a$ | 0.018<br>(0.010, 0.027) | 0.004 | 0.063<br>(0.044, 0.082) | 0.009 |
| $T_{min}$ (℃) | -21.15<br>(-45.65, 3.36) | 11.71 | -7.14<br>(-23.50, 9.21) | 7.81 |
| **Model evaluation** | | | | |
| RSS | 2.5256 | | 2.3557 | |
| MSE | 0.1329 | | 0.1240 | |
| RMSE | 0.3646 | | 0.3521 | |
| AIC | 19.1167 | | 17.6537 | |
| AICc | 15.7833 | | 14.3204 | |
| BIC | 21.2057 | | 19.7427 | |

* Est.: Estimation; SE: Standard error.

** 95%CI: lower and upper 95% confidence intervals.

Tab.8 Specific maximum growth rate fitting results of the full temperature secondary model in Microrisk Lab

| | Square-root model | | Huang square-root model | | Cardinal parameter model | |
|---|---|---|---|---|---|---|
| **Parameter estimation*** | | | | | | |
| Parameters | Est.<br>(95% CI) ** | SE | Est.<br>(95% CI) ** | SE | Est.<br>(95% CI) ** | SE |
| $a$ | 0.043<br>(0.036, 0.051) | 0.004 | 0.087<br>(0.077, 0.098) | 0.005 | -<br>- | - |
| $b$ | 0.15<br>(0.09, 0.20) | 0.02 | 0.79<br>(0.60, 0.99) | 0.09 | -<br>- | - |
| $T_{min}$ (℃) | 4.18<br>(0.97, 7.39) | 1.52 | 3.63<br>(-0.75, 8.02) | 2.08 | 5.64<br>(3.09, 8.19) | 1.21 |
| $T_{max}$ (℃) | 51.91<br>(50.57, 53.26) | 0.64 | 47.49<br>(47.34, 47.63) | 0.07 | 49.60<br>(48.89, 50.32) | 0.339 |
| $\mu_{opt}$ (1/h) | -<br>- | - | -<br>- | - | 1.620<br>(1.558, 1.682) | 0.029 |
| $T_{opt}$ (℃) | -<br>- | - | -<br>- | - | 39.73<br>(38.93, 40.53) | 0.38 |
| **Model evaluation** | | | | | | |
| RSS | 0.0999 | | 0.3881 | | 0.0816 | |
| MSE | 0.0059 | | 0.0228 | | 0.0048 | |
| RMSE | 0.0767 | | 0.1511 | | 0.0693 | |
| AIC | -44.7134 | | -16.2161 | | -48.9750 | |
| AICc | -50.2134 | | -21.7161 | | -54.4750 | |
| BIC | -40.5354 | | -12.0380 | | -44.7969 | |

* Est.: Estimation; SE: Standard error.

** 95%CI: lower and upper 95% confidence intervals.

#### Practical example Ⅳ - Two flora competition growth fitting

(1) In this case, a study on the competition growth of *Escherichia coli* O157:H7 in ground beef and the background microflora (Vimont et al., 2006) was cited for fitting. Note that the dataset was obtain from the R package '*nlsMicrobio*' (Baty & Delignette-Muller, 2015).

(2) Choose the ⑥ section of the 'Competition Models' (see Fig.2), and the setting panel of the competition growth model will show up. The steps of importing data and obtaining results are similar to the isothermal growth fitting (see Fig.4 to Fig.7). While the counting data of the two flora should be input together in the 'Data' box (Fig.8).

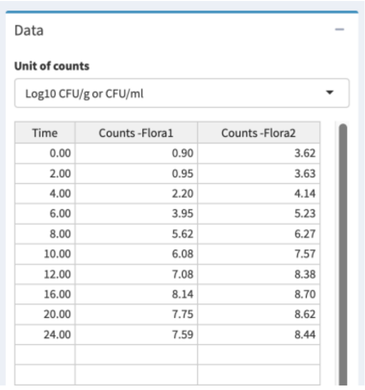
Data

Unit of counts

Log10 CFU/g or CFU/ml

| Time | Counts -Flora1 | Counts -Flora2 |
|---|---|---|
| 0.00 | 0.90 | 3.62 |
| 2.00 | 0.95 | 3.63 |
| 4.00 | 2.20 | 4.14 |
| 6.00 | 3.95 | 5.23 |
| 8.00 | 5.62 | 6.27 |
| 10.00 | 6.08 | 7.57 |
| 12.00 | 7.08 | 8.38 |
| 16.00 | 8.14 | 8.70 |
| 20.00 | 7.75 | 8.62 |
| 24.00 | 7.59 | 8.44 |

Fig.8 Box for the competition growth data input.

(3) For this case, the estimated result of parameters by using different temperature secondary model are listed in Tab.9.

Tab.9 Two flora growth fitting results of the competition model in Microrisk Lab

| | **Jameson-No lag Buchanan model** | | **Jameson-Buchanan model** | |
|---|---|---|---|---|
| **Parameter estimation*** | | | | |
| Parameters | Est. (95% CI) ** | SE | Est. (95% CI) ** | SE |
| $\mu_{max}$ (1/h) - Flora 1 | 1.323 (1.150, 1.467) | 0.081 | 1.566 (1.360, 1.771) | 0.10 |
| $\mu_{max}$ (1/h) - Flora 2 | 1.001 (0.842, 1.160) | 0.075 | 1.323 (1.075, 1.571) | 0.115 |
| $y_0$ (log10 CFU/g) - Flora 1 | 0.38 (-0.21, 0.96) | 0.27 | 0.90 (0.26, 1.55) | 0.30 |
| $y_0$ (log10 CFU/g) - Flora 2 | 2.94 (2.37, 3.52) | 0.27 | 3.63 (3.17, 4.08) | 0.21 |
| $t_{lag}$ (h) - Flora 1 | | | 1.80 (0.60, 3.00) | 0.55 |
| $t_{lag}$ (h) - Flora 2 | | | 3.20 (1.91, 4.48) | 0.59 |
| $t_{max}$ (h) | 12.97 (11.81, 14.13) | 0.54 | 11.73 (10.87, 12.58) | 0.40 |
| **Model evaluation** | | | | |
| RSS | 2.5373 | | 1.1581 | |
| MSE | 0.1692 | | 0.0891 | |
| RMSE | 0.4113 | | 0.2985 | |
| AIC | 25.4652 | | 13.7782 | |
| AICc | 19.7509 | | 9.1115 | |
| BIC | 30.4439 | | 20.7483 | |

* Est.: Estimation; SE: Standard error.

** 95%CI: lower and upper 95% confidence intervals.

#### Practical example V – Non-isothermal growth fitting

(1) In this case, a study on the non-isothermal growth of *L. monocytogenes* in ready-to-eat braised beef was introduced for fitting two dynamic growth models.

(2) Choose the ⑦ 'Growth' in the section of the 'Non-isothermal Models' (see Fig.2), and the setting panel of temperature secondary model will show up.

(3) Both time-temperature profile and the bacterial counting data are required in the non-isothermal modeling (Fig. 9A and Fig. 9B). Meanwhile, the initial guess of the model parameter was also necessary to conduct the regression (Fig. 9C). User can also change the time-step of the regression (default is 0.1). Then the estimated results should be listed if the converge is successful.

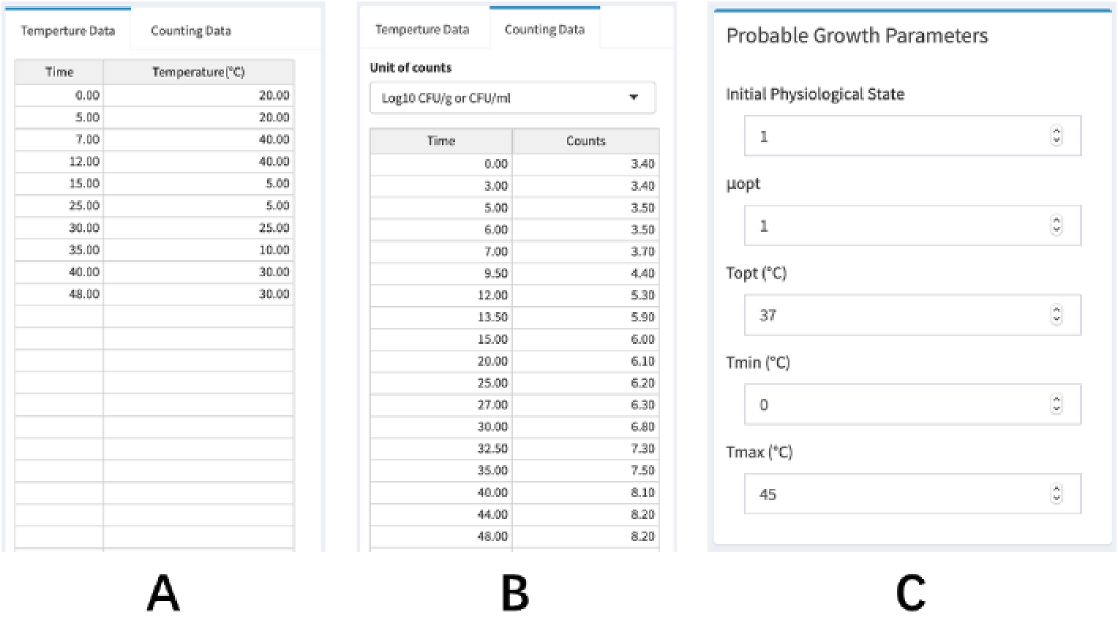

Fig.9 Box for (A) temperature profile and (B) input; and (B) the setting of initial guesses.

(4) For this case, the estimated result of parameters by using two non-isothermal growth model are listed in Tab.10.

Tab.10 Non-isothermal growth fitting results of the dynamic model in Microrisk Lab

| **Baranyi-Cardinal parameter model** | | | | **Huang-Cardinal parameter model** | | | |
|---|---|---|---|---|---|---|---|
| **Parameter estimation*** | | | | | | | |
| Parameters | Int. | Est. (95% CI) ** | SE | Parameters | Int. | Est. (95% CI) ** | SE |
| $y_0$ (log10 CFU/g) | - | 3.39 (3.36, 3.43) | 0.02 | $y_0$ (log10 CFU/g) | - | 3.45 (3.41, 3.50) | 0.02 |
| $y_{max}$ (log10 CFU/g) | - | 8.21 (8.18, 8.25) | 0.02 | $y_{max}$ (log10 CFU/g) | - | 8.21 (8.16, 8.27) | 0.03 |
| $\mu_{opt}$ (1/h) | 1 | 1.065 (0.854, 1.276) | 0.096 | $\mu_{opt}$ (-h) | 1 | 1.242 (0.825, 1.659) | 0.187 |
| $T_{opt}$ (℃) | 37 | 36.4 (35.4, 37.5) | 0.5 | $T_{opt}$ (℃) | 37 | 38 (33.5, 42.4) | 2 |
| $T_{min}$ (℃) | 0 | -1.1 (-2.6, 0.5) | 0.7 | $T_{min}$ (℃) | 0 | -2.8 (-7.4, -1.8) | 2.1 |
| $T_{max}$ (℃) | 45 | 42.4 (38.4, 46.4) | 1.8 | $T_{max}$ (℃) | 45 | 40.3 (38.7, 41.9) | 0.7 |
| $q_0$ | 1 | 0.0244 (0.0167, 0.0321) | 0.0035 | $A$ | 1 | 1.91 (1.84, 1.99) | 0.04 |
| | | | | $m$ | 1 | 0.33 (0.17, 0.48) | 0.07 |
| **Step size (h)** | | | | 0.1 | | | |
| **Model evaluation** | | | | | | | |
| RSS | 0.0071 | | | | 0.0155 | | |
| MSE | 0.0006 | | | | 0.0016 | | |
| RMSE | 0.0253 | | | | 0.0394 | | |
| AIC | -46.0602 | | | | -29.8674 | | |
| AICc | -48.8602 | | | | -29.8674 | | |
| BIC | -39.8276 | | | | -22.7444 | | |

* Int.: Initial guess; Est.: Estimation; SE: Standard error.

** 95%CI: lower and upper 95% confidence intervals.

#### Practical example VI – Non-isothermal inactivation fitting

(1) In this case, a study on the non-isothermal inactivation of *Bacillus sporothermodurans* IC4 spores under dynamic heating conditions (Garre et al, 2018) was cited for fitting two dynamic inactivation models.

(2) Choose the ⑧ 'Inactivation' in the section of the 'Non-isothermal Models' (see Fig.2), and the setting panel of temperature secondary model will show up. The steps of importing data and obtaining results are similar to the non-isothermal growth fitting (see Fig. 9).

(3) For this case, the estimated result of parameters by using two non-isothermal inactivation model are listed in Tab.11.

Tab.11 Non-isothermal inactivation fitting results of the dynamic model in Microrisk Lab

| **Dynmaic Bigelow model** | | | | **Dynamic Weibull model** | | | |
|---|---|---|---|---|---|---|---|
| **Parameter estimation*** | | | | | | | |
| Parameters | Int. | Est. (95% CI) ** | SE | Parameters | Int. | Est. (95% CI) ** | SE |
| $T_{ref}$ (°C) | 130 | - - | - | $T_{ref}$ (°C) | 130 | - - | - |
| $y_0$ (log10 CFU/g) | - | 5.78 (5.69, 5.87) | 0.04 | $y_0$ (log10 CFU/g) | - | 5.78 (5.69, 5.88) | 0.04 |
| $D_{ref}$ (min) | 2 | 0.18 (0.05, 0.31) | 0.06 | $\delta_{ref}$ | 2.00 | 4.88 (-8.72, 18.48) | 6.42 |
| $z$ (°C) | 6 | 6.67 (4.7, 8.6) | 0.92 | $z$ (°C) | 6.00 | 9.79 (-3.52, 23.09) | 6.28 |
| | | | | $p$ | 1.00 | 1.35 (0.01, 2.69) | 0.63 |
| **Step size (h)** | | | | 0.01 | | | |
| **Model evaluation** | | | | | | | |
| RSS | 0.1737 | | | | 0.1704 | | |
| MSE | 0.0102 | | | | 0.0106 | | |
| RMSE | 0.1011 | | | | 0.1032 | | |
| AIC | -32.1667 | | | | -30.5504 | | |
| AICc | -36.6667 | | | | -35.8837 | | |
| BIC | -29.1795 | | | | -26.5675 | | |

* Int.: Initial guess; Est.: Estimation; SE: Standard error.

** 95%CI: lower and upper 95% confidence intervals.

### 7 Simulation module of Microrisk Lab

The simulation module allows to solve the ①② isothermal and ③④ non-isothermal forward problem in predictive microbiology (Fig.10). There are no limitations in the condition setting. Users may simulate the bacterial growth or inactivation with the prior knowledge on the kinetic parameter and growth/ death boundary. Moreover, both deterministic and stochastic models are provided in the isothermal simulation.

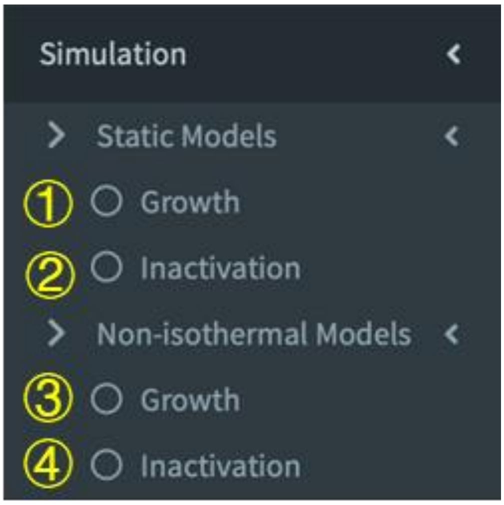

Fig.10 Different sections of model fitting in the simulation module.

##### Practical example VII- Stochastic growth simulation

(1) The condition setting of the growth simulation is adopted from the stochastic growth of *Salmonella* Typhimurium individual cells researched by Koutsoumanis and Lianou (2013). Tab.12 lists the setting for simulation. The Buchanan model is chosen as the growth model for individual cells. A 10,000 times iteration was realized based on the simple sampling method for Monte-Carlo simulation.

Tab.12 Stochastic growth simulation settings for Microrisk Lab

| **Parameters** | **Microrisk Lab** | |
|---|---|---|
| $y_0$ ($\log_{10}$ CFU/g) | Distribution | Normal |
| | Mean | 0 |
| | Standard deviation | 0 |
| $y_{max}$ ($\log_{10}$ CFU/g) | Distribution | Normal |
| | Mean | 8 |
| | Standard deviation | 0 |
| $t_{lag}$ | Distribution | LogNormal |
| | Mean | 3.355 |
| | Standard deviation | 0.896 |
| | Shift | -1.628 |
| $\mu_{max}$ | Distribution | Logistic |
| | Mean | 0.754 |
| | Standard deviation | 0.024 |
| $t$ | Distribution | Uniform |
| | Maximum | 0 |
| | Minimum | 8 |
| **Model** | Buchanan model | |
| **Iteration times** | 10,000 | |

(2) Choose ① the 'Growth' section of the 'Isothermal Models' in the 'Simulation' module, and ② choose 'Stochastic' model type in the setting panel (Fig.11). Then set the ③ 'Iteration time' and ④ 'Model' to '10,000' and 'Buchanan model', respectively.

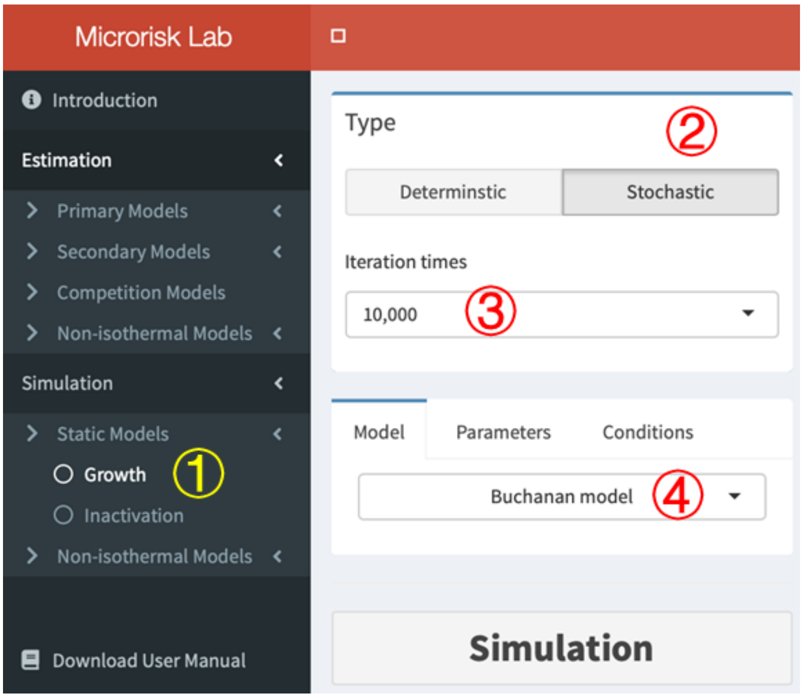

Fig.11 Layout of a section of the 'Simulation' module.

(3) Switch to the ① ‘Parameters’ tab to determine the setting of the (distribution of) ② $y_{max}$, ③ $t_{lag}$, and ④ $\mu_{max}$ according to Tab.12 (Fig.12).

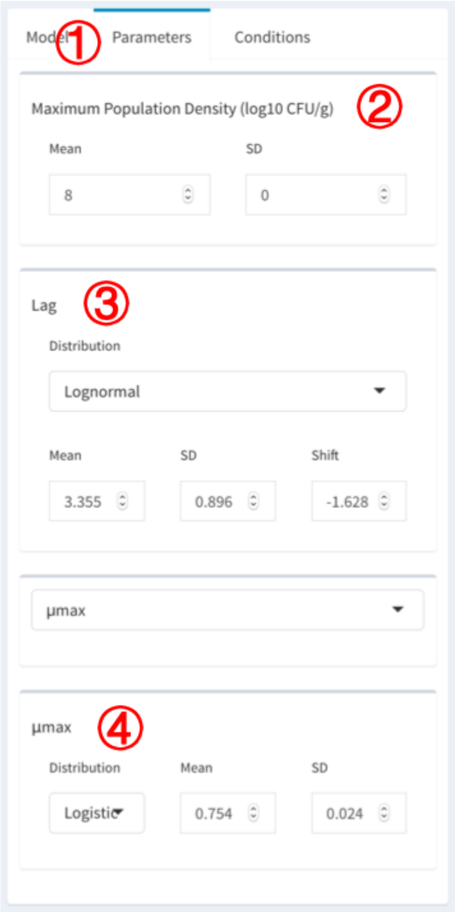

Fig.12 Box for kinetic parameter setting.

(4) Switch to the ① ‘Conditions’ tab to determine the setting of the (distribution of) ② $y_0$, ③ $t$ according to Tab.12 (Fig.13).

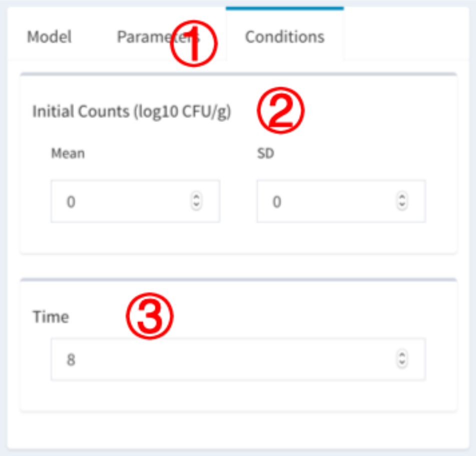

Fig.13 Box for condition setting.

(5) Click the ① 'Simulation' button. After a necessary loading time, if no contradiction in the setting, the ② simulated curve/point and ③ predicted result will show in the result panel (Fig.14A). Otherwise, different popup messages will appear for the simulation warning (Fig.14B-D). In these cases, please check the setting of kinetic parameters and the condition of the simulated environment.

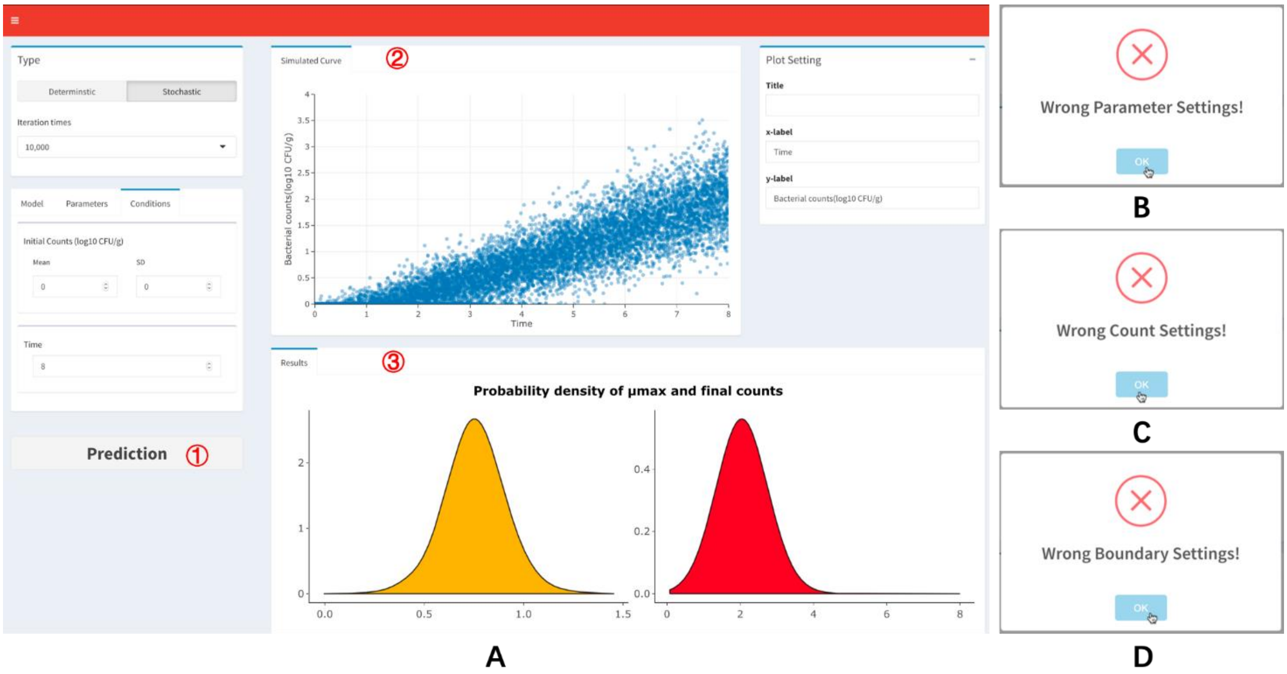

Fig.14 Layout of the interface after simulation.

(6) The stochastic growth simulation can be viewed on ① the interactive plot, which is also adjustable and downloadable (Fig.15A). The distribution of ② the estimated $\mu_{max}$ and ③ final bacterial concentration ($y_{final}$) , as well as ④ the estimated mean value and standard deviation will

be presented and listed (Fig.15B). The sensitivity analysis on model parameters is realized by calculating the Pearson correlation between different factors and the bacterial counts. Here, according to ⑤ the correlation plot, the duration of growth time is the most sensitive parameter for the bacterial counts during the stochastic growth of *S.* Typhimurium single cell (Fig.15B).

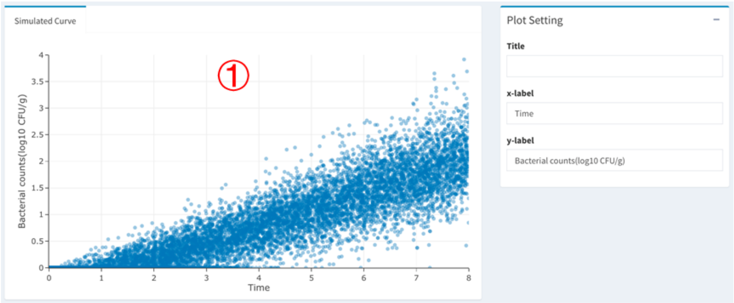

A

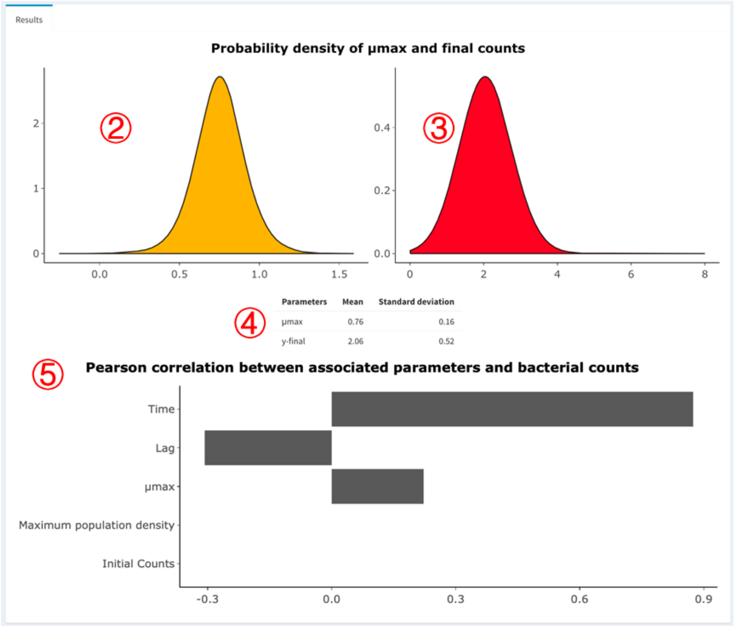

B

Fig.15 The result of stochastic simulation.

(7) Note that, in the section of 'Non-isothermal Models' of the simulation module, only deterministic model was provided in this version.

### 8 Predictive models integrated in Microrisk Lab

Microrisk Lab consists of 11 isothermal growth models (Tab.13), 9 inactivation models (Tab.14),10 secondary models (Tab.15), 2 competition growth models (Tab.16), and 4 non-isothermal models (Tab.17) for estimation or simulation work.

Tab.13. Explicit equations for growth included in Microrisk Lab

| Name | Formula |
|---|---|
| **Complete model** | |
| Gompertz model [1] | $Y(t) = Y_0 + (Y_{max} - Y_0)exp\left\{-\exp\left[\frac{2.71\mu_{max}(t_{lag}-t)}{Y_{max}-Y_0} + 1\right]\right\}$ |
| Baranyi model [2] | $\begin{cases} Y(t) = Y_0 + \mu_{max}A(t) - \ln\left[1 + \frac{\exp(\mu_{max}A(t))-1}{\exp(Y_{max}-Y_0)}\right] \\ A(t) = t + \frac{1}{\mu_{max}}\left[\ln\exp(-\mu_{max}t) + \exp(-\mu_{max}t_{lag}) - \exp(-\mu_{max}t - \mu_{max}t_{lag})\right] \end{cases}$ |
| Buchanan model [3] | $\begin{cases} y(t) = y_0,\ t < t_{lag} \\ y(t) = y_0 + \frac{\mu_{max}}{\ln 10}(t - t_{lag}),\ t_{lag} \le t < t_{max} \\ y(t) = y_{max},\ \ t \ge t_{max} \end{cases}$ |
| Lag-logistic model [4] | $\begin{cases} Y(t) = Y_0,\ t < t_{lag} \\ Y(t) = Y_{max} - \ln\{1 + [\exp(Y_{max} - Y_0) - 1]\exp[-\mu_{max}(t - t_{lag})]\},\ t \ge t_{lag} \end{cases}$ |
| Huang model [5] | $\begin{cases} Y(t) = Y_0 + Y_{max} - \ln\{\exp(Y_0) + [\exp(Y_{max}) - \exp(Y_0)]\exp(-\mu_{max}B(t))\} \\ B(t) = t + \frac{1}{4}\ln\frac{1+\exp[-4(t-t_{lag})]}{1-\exp(4t_{lag})} \end{cases}$ |
| **No lag model** | |
| Logistic model [6] | $Y(t) = Y_0 + Y_{max} - \ln\{\exp(Y_0) + [\exp(Y_{max}) - \exp(Y_0)]\exp(-\mu_{max}t)\}$ |
| Buchanan model [7] | $\begin{cases} y(t) = y_0 + \frac{\mu_{max}}{\ln 10}t,\ t < t_{max} \\ y(t) = y_{max},\ t \ge t_{max} \end{cases}$ |
| **Reduced model** | |
| Baranyi model [8] | $Y(t) = Y_0 + \mu_{max}t + \ln[\exp(-\mu_{max}t) + \exp(-\mu_{max}t_{lag}) - \exp(-\mu_{max}t - \mu_{max}t_{lag})]$ |
| Buchanan model [9] | $\begin{cases} y(t) = y_0,\ t < t_{lag} \\ y(t) = y_0 + \frac{\mu_{max}}{\ln 10}(t - t_{lag}),\ t \ge t_{lag} \end{cases}$ |
| Huang model [10] | $Y(t) = Y_0 + \mu_{max}t + \frac{1}{4}\mu_{max}\ln\frac{1+\exp[-4(t-t_{lag})]}{1-\exp(4t_{lag})}$ |
| **Linear model** | |
| Linear model | $Y(t) = Y_0 + \mu_{max}t$ |

[1]Zwietering et al., 1990; [2/8] Baranyi and Roberts, 1995; [3/7/9] Buchanan et al., 1997; [4] Rosso et al., 1996; [5/6/10] Huang, 2008.

Tab.14. Explicit equations for inactivtion included in Microrisk Lab

| Name | Formula |
|---|---|
| **Complete model** | |
| Completed Geeraerd model [1] | $y(t) = y_{res} + \log_{10}\left[\frac{(10^{y_0-y_{res}}-1)\exp(k_{max}S_l)}{\exp(k_{max}t)+\exp(k_{max}S_l)-1} + 1\right]$ |
| Three-phase model [2] | $\begin{cases} y(t) = y_0,\ t < S_l \\ y(t) = y_0 + \frac{k_{max}}{\ln 10}(t - S_l),\ S_l \le t < S_t \\ y(t) = y_{res},\quad t \ge S_t \end{cases}$ |
| Weibull-tail model [3] | $y(t) = y_{res} + \log_{10}\left[(10^{y_0-y_{res}} - 1)\,10^{-\left(\frac{t}{\delta}\right)^p} + 1\right]$ |
| **No shoulder model** | |
| No shoulder Geeraerd model [4] | $y(t) = y_{res} + \log_{10}\{(10^{y_0-y_{res}} - 1)\exp(k_{max}t) + 1\}$ |
| No shoulder two-phase model [5] | $\begin{cases} y(t) = y_0 + \frac{k_{max}}{\ln 10}t, \quad t < S_t \\ y(t) = y_{res},\quad t \ge S_t \end{cases}$ |
| **No tail model** | |
| No tail Geeraerd model [6] | $y(t) = y_0 + \frac{k_{max}t}{\ln 10} + \log_{10}\left\{\frac{\exp(k_{max}S_l)}{1+[\exp(k_{max}S_l)-1]\exp(k_{max}t)}\right\}$ |
| No tail two-phase model [7] | $\begin{cases} y(t) = y_0,\ t < S_l \\ y(t) = y_0 + \frac{k_{max}}{\ln 10}(t - S_l),\quad t \ge S_l \end{cases}$ |
| Weibull model [8] | $y(t) = y_0 - \left(\frac{t}{\delta}\right)^p$ |
| **Linear model** | |
| Bigelow model [9] | $y(t) = y_0 - \frac{t}{D}$ |

[1/4/6] Geeraerd et al., 2000; [2/5/7] Buchanan and Golden, 1995; [3] Albert and Mafart, 2005; [8] van Boekel, 2002; [9] Bigelow, 1921.

Tab.15. Secondary models for $\mu_{max}$ included in Microrisk Lab

| Name | Formula |
|---|---|
| **Temperature models** | |
| Suboptimal square-root model [1] | $\mu_{max} = [a(T - T_{min})]^2$ |
| Full square-root model [2] | $\mu_{max} = \langle a(T - T_{min})\{1 - \exp[b(T - T_{max})]\}\rangle^2$ |
| Suboptimal Huang square-root model [3] | $\mu_{max} = [a(T - T_{min})^{0.75}]^2$ |
| Full Huang square-root model [4] | $\mu_{max} = \langle a(T - T_{min})^{0.75}\{1 - \exp[b(T - T_{max})]\}\rangle^2$ |
| Cardinal parameter model [5] | $\mu_{max} = \frac{\mu_{opt}(T-T_{max})(T-T_{min})^2}{[(T_{opt}-T_{min})(T-T_{opt})-(T_{opt}-T_{max})(T_{opt}+T_{min}-2T)](T_{opt}-T_{min})}$ |
| **pH models** | |
| Cardinal 3-parameter model [6] | $\mu_{max} = \frac{\mu_{opt}(pH-pH_{min})[pH-(2pH_{opt}-pH_{min})]}{(pH-pH_{min})[pH-(2pH_{opt}-pH_{min})]-(pH-pH_{opt})^2}$ |
| Cardinal 4-parameter model [7] | $\mu_{max} = \frac{\mu_{opt}(pH-pH_{min})(pH-pH_{max})}{(pH-pH_{min})(pH-pH_{max})-(pH-pH_{opt})^2}$ |
| Quasi-mechanistic model [8] | $\mu_{max} = \mu_{opt}(1 - 10^{pH_{min}-pH})$ |
| **Water activity models** | |
| Cardinal 2-parameter model [9] | $\mu_{max} = \frac{\mu_{opt}(aw-aw_{min})^2}{(1-aw_{min})^2}$ |
| Cardinal 3-parameter model [10] | $\mu_{max} = \frac{\mu_{opt}(aw-1)(aw-aw_{min})^2}{(aw_{opt}-aw_{min})[(aw_{opt}-aw_{min})(aw-aw_{opt})-(aw_{opt}-1)(aw_{opt}+aw_{min}-2aw)]}$ |

[1/2] Ratkowsky et al., 1983; [3/4] Huang and Hwang, 2011; [5] Rosso et al, 1993; [6/7] Rosso et al, 1995;[8] Presser et al. 1997; [9/10] Rosso and Robinson, 2001

Tab.16. Two flora competition growth models included in Microrisk Lab

| Name | Formula |
|---|---|
| Jameson - No lag Buchanan model [1] | $\begin{cases} y_1(t) = \begin{cases} y_1 + \frac{\mu_{max1}}{\ln 10} t, \ t < t_{max} \\ y_1 + \frac{\mu_{max1}}{\ln 10} t_{max}, \ t \geq t_{max} \end{cases} \\ y_2(t) = \begin{cases} y_2 + \frac{\mu_{max2}}{\ln 10} t, \ t < t_{max} \\ y_2 + \frac{\mu_{max2}}{\ln 10} t_{max}, \ t \geq t_{max} \end{cases} \end{cases}$ |
| Jameson - Buchanan model [2] | $\begin{cases} y_1(t) = \begin{cases} y_1, \ t < t_{lag1} \\ y_1 + \frac{\mu_{max1}}{\ln 10}(t - t_{lag1}), \ t_{lag1} \leq t < t_{max} \\ y_1 + \frac{\mu_{max1}}{\ln 10}(t_{max} - t_{lag1}), \ t \geq t_{max} \end{cases} \\ y_2(t) = \begin{cases} y_2, \ t < t_{lag2} \\ y_2 + \frac{\mu_{max2}}{\ln 10}(t - t_{lag2}), \ t_{lag2} \leq t < t_{max} \\ y_2 + \frac{\mu_{max2}}{\ln 10}(t_{max} - t_{lag2}), \ t \geq t_{max} \end{cases} \end{cases}$ |

* The inferior number 1 or 2 in competition growth models represent the flora type.

[1/2] Vimont et al., 2006

Tab.17. Ordinary differential equations for growth/ inactivtion included in Microrisk Lab

| Name | Formula |
|---|---|
| **Non-isothermal growth models** | |
| Baranyi - Cardinal parameter model [1] | $\begin{cases} \frac{dY}{dt} = \mu_{max}\left[\frac{1}{1+\exp(-Q)}\right][1 - \exp(Y - Y_{max})] \\ \frac{dQ}{dt} = \mu_{max} \\ Q = \ln\frac{q}{1-q} \\ Y(0) = Y_0 \\ q(0) = q_0 \\ \mu_{max} = \frac{\mu_{opt}(T-T_{max})(T-T_{min})^2}{[(T_{opt}-T_{min})(T-T_{opt})-(T_{opt}-T_{max})(T_{opt}+T_{min}-2T)](T_{opt}-T_{min})} \end{cases}$ |
| Huang - Cardinal parameter model [2/3] | $\begin{cases} \frac{dY}{dt} = \mu_{max}\left[\frac{1}{1+\exp(-4(t-t_{lag}))}\right][1 - \exp(Y - Y_{max})] \\ t_{lag} = \frac{\exp(A)}{{\mu_{max}}^m} \\ Y(0) = Y_0 \\ \mu_{max} = \frac{\mu_{opt}(T-T_{max})(T-T_{min})^2}{[(T_{opt}-T_{min})(T-T_{opt})-(T_{opt}-T_{max})(T_{opt}+T_{min}-2T)](T_{opt}-T_{min})} \end{cases}$ |
| **Non-isothermal inactivation model** | |
| Dynamic Weibull model [4] | $\frac{dy}{dt} = -p\left(\frac{10^{\frac{T-T_{ref}}{z}}}{\delta_{ref}}\right)^p t^{p-1}, y(0) = y_0$ |
| Dynamic Bigelow model [5] | $\frac{dy}{dt} = -\frac{1}{D_{ref}} 10^{\frac{T-T_{ref}}{z}}, y(0) = y_0$ |

[1/2/3] Huang, 2017; [4] Mafart et al, 2002;[5] Van Impe et al., 1992.

### 9 Statistical indicators in Microrisk Lab

To evaluate and compare the goodness of fit, the statistical indicator of residual sum of squares (RSS, Eq.1), mean square error (MSE, Eq.2), root mean square error (RMSE, Eq.3), regular Akaike information criterion (AIC, Eq.4, Akaike, 1974), modified AIC (AICc, Eq.5, Burnham & Anderson, 2003) and Bayesian information criterions (BIC, Eq.6, Schwarz, 1978) are provided such in the 'Model Evaluation' tab for all regression analyses. The coefficient of determination (R-square R2, Eq.7) and adjusted coefficient of determination (Adjusted $R^2$, Eq.8) were provided only for linear models.

$$\mathrm{RSS} = \sum_{i=1}^{n}(y_i - \hat{y}_i)^2 \qquad \text{Eq.1}$$

$$\mathrm{MSE} = \frac{\mathrm{RSS}}{n-k} \qquad \text{Eq.2}$$

$$\mathrm{RMSE} = \sqrt{\mathrm{MSE}} \qquad \text{Eq.3}$$

$$\mathrm{AIC} = -2\log\left(\hat{\theta}\right) + 2k \qquad \text{Eq.4}$$

$$\mathrm{AIC_c} = \mathrm{AIC} + \frac{2k(k+1)}{n-k-1} \qquad \text{Eq.5}$$

$$\mathrm{BIC} = -2\log\left(\hat{\theta}\right) + k\ln(n) \qquad \text{Eq.6}$$

$$\mathrm{R}^2 = \frac{\sum_{i=1}^{n}\left(\hat{y}_i - \frac{1}{n}\sum_{i=1}^{n} y_i\right)^2}{\sum_{i=1}^{n}\left(y_i - \frac{1}{n}\sum_{i=1}^{n} y_i\right)^2} \qquad \text{Eq.7}$$

$$\text{Adjusted } \mathrm{R}^2 = 1 - (1 - \mathrm{R}^2)\frac{n-1}{n-k-1} \qquad \text{Eq.8}$$

where $y_i$ is the i th value of the observation; $\hat{y}_i$ is the i th value of the prediction; $k$ is the number of parameters; and $n$ is the number of sample data; $\log\left(\hat{\theta}\right)$ is the numerical value of the log-likelihood for the fitted model (the probability of the data given a model in the model).

### Reference

Akaike, H. (1974). A new look at the statistical model identification. *IEEE Transactions on Automatic Control*, *19*(6), 716–723.

Albert, I., & Mafart, P. (2005). A modified Weibull model for bacterial inactivation. *International Journal of Food Microbiology*, *100*(1-3), 197–211.

Attali, D., Cheng, J. & Edwards, T. (2019). *shinyalert*: Easily Create Pretty Popup Messages (Modals) in 'Shiny' *R package version 1.0*. Available at: www.r-project.org.

Baranyi, J., & Roberts, T. A. (1995). Mathematics of predictive food microbiology. *International Journal of Food Microbiology*, *26*(2), 199–218.

Baty, F., & Delignette-Muller, M.-L. (2015). *nlsMicrobio*: Nonlinear regression in predictive microbiology. *R package version 0.0-1*. Available at: www.r-project.org.

Bigelow, W. D. (1921). The logarithmic nature of thermal death time curves. *Journal of Infectious Diseases*, *29*(5), 528–536.

Buchanan, R. L., & Golden, M. H. (1995). Model for the non-thermal inactivation of *Listeria monocytogenes* in a reduced oxygen environment. *Food Microbiology*, *12*, 203–212.

Buchanan, R. L., Whiting, R. C., & Damert, W. C. (1997). When is simple good enough: a comparison of the Gompertz, Baranyi, and three-phase linear models for fitting bacterial growth curves. *Food Microbiology*, *14*(4), 313–326.

Burnham, K. P., & Anderson, D. R. (2003). *Model Selection and Multimodel Inference*. Springer Science & Business Media.

Chang, W., Cheng, J., Allaire, J., Xie, Y., & McPherson, J. (2019). *shiny*: web application framework for R. *R package version 1.0.5*. Available at: www.r-project.org.

Chang, W., & Borges Ribeiro, B. (2019). *shinydashboard*: create dashboards with'Shiny'. *R package version 0.7.1*. Available at: www.r-project.org.

Garre, A., Clemente-Carazo, M., Fernández, P. S., Lindqvist, R., & Egea, J. A. (2018). Bioinactivation FE: A free web application for modelling static and dynamic microbial inactivation. *Food Research International*, *112*, 353–360. http://doi.org/10.1016/j.foodres.2018.06.057

Geeraerd, A. H., Herremans, C. H., & Van Impe, J. F. (2000). Structural model requirements to describe microbial inactivation during a mild heat treatment. *International Journal of Food Microbiology*, *59*(3), 185–209.

Hamner, B., Frasco, M., & LeDell, E. (2018). *Metrics*: Evaluation metrics for machine learning. *R package version 0.1.4*. Available at: www.r-project.org.

Huang, L. (2008). Growth kinetics of *Listeria monocytogenes* in broth and beef Frankfurters—Determination of lag phase duration and exponential growth rate under static conditions. *Journal of Food Science*, *73*(5), E235–E242.

Huang, L. (2014). IPMP 2013--a comprehensive data analysis tool for predictive microbiology. *International Journal of Food Microbiology*, *171*, 100–107.

Huang, L. (2017a). Dynamic identification of growth and survival kinetic parameters of microorganisms in foods. *Current Opinion in Food Science*, *14*, 85–92.

Huang, L., Hwang, C.-A., & Phillips, J. (2011). Evaluating the effect of temperature on microbial growth rate-The Ratkowsky and a Bělehrádek-type models. *Journal of Food Science*, *76*(8), M547–M557.

Koutsoumanis, K. P., & Lianou, A. (2013). Stochasticity in colonial growth dynamics of individual bacterial cells. *Applied and Environmental Microbiology*, *79*(7), 2294–2301.

Mafart, P., Couvert, O., Gaillard, S., & Leguérinel, I. (2002). On calculating sterility in thermal preservation methods: application of the Weibull frequency distribution model. *International Journal of Food Microbiology*, *72*(1-2), 107–113.

Owen, J., Allen, J., Hocking, T., Xie, Y., Martoglio, E., Ger, I., Marcin, W. et al. (2019). *rhandsontable*: Interface to the 'Handsontable.js' Library. *R package version 0.3.7.* Available at: www.r-project.org.

Perrier, V., Meyer, F., & Granjon, D. (2019). *shiny*: web application framework for R. *R package version 0.4.8*. Available at: www.r-project.org.

Pouillot, R., Delignette-Muller, M., & Denis, J. (2017). *mc2d*: tools for two-dimensional Monte-Carlo simulations. *R package version 0.1-18*. Available at: www.r-project.org.

Presser, K. A., Ratkowsky, D. A., & Ross, T. (1997). Modelling the growth rate of *Escherichia coli* as a function of pH and lactic acid concentration. *Applied and Environmental Microbiology*, *63*(6), 2355–2360.

Ratkowsky, D. A., Lowry, R. K., McMeekin, T. A., Stokes, A. N., & Chandler, R. E. (1983). Model for bacterial culture growth rate throughout the entire biokinetic temperature range. *Journal of Bacteriology*, *154*(3), 1222–1226.

Rosso, L., Bajard, S., Flandrois, J. P., Lahellec, C., Fournaud, J., & Veit, P. (1996). Differential growth of *Listeria monocytogenes* at 4 and 8°C: Consequences for the Shelf Life of Chilled Products. *Journal of Food Protection*, *59*(9), 944–949.

Rosso, L., Lobry, J. R., Bajard, S., & Flandrois, J. P. (1995). Convenient model to describe the combined effects of temperature and pH on microbial growth. *Applied and Environmental Microbiology*, *61*(2), 610–616.

Rosso, L., Lobry, J. R., & Flandrois, J. P. (1993). An unexpected correlation between cardinal temperatures of microbial growth highlighted by a new model. *Journal of Theoretical Biology*, *162*(4), 447–463.

Rosso, L., & Robinson, T. P. (2001). A cardinal model to describe the effect of water activity on the growth of moulds. *International Journal of Food Microbiology*, *63*(3), 265–273.

Schwarz, G. (1978). Estimating the dimension of a model. *The Annals of Statistics*, *6*(2), 461–464.

Sievert, C., Parmer, C., Hocking, T., Chamberlain, S., Ram, K., Corvellec, M., & Despouy, P. (2019). *plotly*: Create Interactive Web Graphics via'plotly. js'. *R package version 4.9.0.* Available at: www.r-project.org.

van Boekel, M. A. J. S. (2002). On the use of the Weibull model to describe thermal inactivation of microbial vegetative cells. *International Journal of Food Microbiology*, *74*(1-2), 139–159.

Van Impe, J. F., Nicolaï, B. M., Martens, T., De Baerdemaeker, J., & Vandewalle, J. (1992). Dynamic mathematical model to predict microbial growth and inactivation during food processing. *Applied and Environmental Microbiology*, *58*(9), 2901–2909.

Vimont, A., Vernozy-Rozand, C., Montet, M. P., Lazizzera, C., Bavai, C., & Delignette-Muller, M. L. (2006). Modeling and predicting the simultaneous growth of *Escherichia coli* O157:H7 and ground beef background microflora for various enrichment protocols. *Applied and Environmental Microbiology*, *72*(1), 261–268.

Wickham, H., & Chang, W. (2019). *ggplot2*: An implementation of the Grammar of Graphics. *R package version 2.2.1.* Available at: www.r-project.org.

Zwietering, M. H., Jongenburger, I., Rombouts, F. M., & van 't Riet, K. (1990). Modeling of the bacterial growth curve. *Applied and Environmental Microbiology*, *56*(6), 1875–1881.
