## Supplementary material for "Microrisk Lab: an online freeware for predictive microbiology": S.2-Referenced data used in manuscript.pdf

***Case I – Kinetic analysis of* Listera monocytogenes/ innocua *isothermal growth***

| Time (h) | Counts (Log 10 CFU/g) |
|---|---|
| 0 | 3.86 |
| 72 | 4.53 |
| 144 | 5.86 |
| 192 | 7.00 |
| 240 | 7.65 |
| 336 | 9.12 |
| 360 | 9.39 |
| 408 | 9.39 |

Reference ComBASE browser (www.combase.cc, ID: LM127_11)

***Case II – Kinetic analysis of* Salmonella enterica *isothermal inactivation***

| Time (h) | Counts (Log 10 CFU/g) |
|---|---|
| 0.00 | 9.08 |
| 0.17 | 8.95 |
| 0.33 | 8.77 |
| 0.50 | 8.67 |
| 0.67 | 8.23 |
| 0.83 | 7.91 |
| 1.00 | 7.56 |
| 1.17 | 7.39 |
| 1.33 | 7.00 |
| 1.50 | 6.46 |
| 1.67 | 6.07 |
| 1.83 | 5.62 |
| 2.00 | 5.03 |

Reference

Wang, X., Devlieghere, F., Geeraerd, A., & Uyttendaele, M. (2017). Thermal inactivation and sublethal injury kinetics of *Salmonella enterica* and *Listeria monocytogenes* in broth versus agar surface. *International Journal of Food Microbiology*, 243, 70–77. http://doi.org/10.1016/j.ijfoodmicro.2016.12.008

***Case III – Effect of temperature on the specific growth rate of* Salmonella *Typhimurium***

| Temperature (℃) | μmax (1/h) |
|---|---|
| 8 | 0.012 |
| 10 | 0.058 |
| 12 | 0.104 |
| 14 | 0.182 |
| 16 | 0.237 |
| 18 | 0.334 |
| 20 | 0.442 |
| 22 | 0.569 |
| 24 | 0.790 |
| 26 | 0.840 |
| 28 | 1.009 |
| 30 | 1.213 |
| 32 | 1.388 |
| 34 | 1.398 |
| 36 | 1.411 |
| 38 | 1.610 |
| 40 | 1.729 |
| 42 | 1.402 |
| 44 | 1.492 |
| 46 | 1.184 |
| 48 | 0.599 |

Reference

Oscar, T. P. (2002). Development and validation of a tertiary simulation model for predicting the potential growth of *Salmonella* typhimurium on cooked chicken. *International Journal of Food Microbiology*, 76(3), 177–190. http://doi.org/10.1016/s0168-1605(02)00025-9

***Case IV – Kinetic analysis of* L. monocytogenes *non-isothermal growth***

| Time (h) | Temperature (℃) |
|---|---|
| 0 | 20 |
| 5 | 20 |
| 7 | 40 |
| 12 | 40 |
| 15 | 5 |
| 25 | 5 |
| 30 | 25 |
| 35 | 10 |
| 40 | 30 |
| 48 | 30 |

| Time (h) | Counts (Log 10 CFU/g) |
|---|---|
| 0.0 | 3.40 |
| 3.0 | 3.40 |
| 5.0 | 3.50 |
| 6.0 | 3.50 |
| 7.0 | 3.70 |
| 9.5 | 4.40 |
| 12.0 | 5.30 |
| 13.5 | 5.90 |
| 15.0 | 6.00 |
| 20.0 | 6.10 |
| 25.0 | 6.20 |
| 27.0 | 6.30 |
| 30.0 | 6.80 |
| 32.5 | 7.30 |
| 35.0 | 7.50 |
| 40.0 | 8.10 |
| 44.0 | 8.20 |
| 48.0 | 8.20 |

Reference Unpublished data

*Case V – Kinetic analysis of* **Bacillus sporothermodurans** *IC4 non-isothermal inactivation*

| Time (min) | Temperature (℃) | Time (min) | Counts (CFU/g) |
|---|---|---|---|
| 0.00 | 75 | 0.00 | 1070000 |
| 7.69 | 95 | 7.69 | 403500 |
| 15.38 | 115 | 15.38 | 611500 |
| 16.15 | 117 | 16.15 | 702000 |
| 16.92 | 119 | 16.92 | 436500 |
| 17.69 | 121 | 17.69 | 253500 |
| 18.46 | 123 | 18.46 | 154500 |
| 20.46 | 123 | 20.46 | 18150 |
| 22.46 | 123 | 22.46 | 2220 |
| 24.46 | 123 | 24.46 | 140 |
| 0.00 | 75 | 0.00 | 481500 |
| 4.69 | 95 | 4.69 | 536000 |
| 15.38 | 115 | 15.38 | 549500 |
| 16.15 | 117 | 16.15 | 455500 |
| 16.92 | 119 | 16.92 | 407000 |
| 17.69 | 121 | 17.69 | 295500 |
| 18.46 | 123 | 18.46 | 195000 |
| 20.46 | 123 | 20.46 | 16400 |
| 22.46 | 123 | 22.46 | 1385 |
| 24.46 | 123 | 24.46 | 175 |

Reference

Garre, A., Clemente-Carazo, M., Fernández, P. S., Lindqvist, R., & Egea, J. A. (2018). Bioinactivation FE: A free web application for modelling static and dynamic microbial inactivation. Food Research International, 112, 353–360. http://doi.org/10.1016/j.foodres.2018.06.057
